# Upregulation of CD55 complement regulator in distinct PBMC subpopulations of COVID-19 patients is associated with suppression of interferon responses

**DOI:** 10.1101/2022.10.07.510750

**Authors:** M. G. Detsika, M. Sakkou, V. Triantafillidou, D. Konstantopoulos, E. Grigoriou, K. Psarra, E. Jahaj, I Dimopoulou, S. E. Orfanos, A. Tsirogianni, G. Kollias, A. Kotanidou

**Author notes:** **Corresponding author:** Prof Anastasia Kotanidou, MD, PhD., 3 Ploutarchou st, Athens, 10675, Greece., **Email address:**.

## Abstract

Complement activation has been verified in COVID-19 patients by both increased serum levels of complement factors C3a and C5b-9 and increased complement deposition at the tissue levels. Complement regulatory proteins (CRPs) CD55, CD46, CD59 and CR1 act to control complement overactivation and eliminate complement deposition and cell lysis. The aim of the study was to investigate the expression of CRPs in COVID-19 in order to identify potential dysregulated expression patterns of CRPs and address whether these may contribute to disease pathogenesis.

Single cell RNA-sequencing (scRNA-seq) analysis performed on isolated PBMCs revealed an increase of CD55 expression in severe and critical COVID-19 patients compared to healthy controls. This increase was also detected upon integrated subclustering analysis of the monocyte, T cell and B cell populations. Flow cytometric analysis verified the distinct pattern of upregulated CD55 expression in monocyte and T cell sub populations of severe COVID-19 patients. This upregulation was associated with decreased expression of interferon stimulated genes (ISGs) in patients with severe COVID-19 suggesting a potential suppressor effect of CD55 on interferon responses. The present study identifies a COVID-19 specific CD55 expression pattern in PBMC subpopulations that coincides with reduced interferon responses thus indicating that the complement regulator CD55 may contribute to COVID-19 pathogenesis.

## Introduction

The complement system acts as an immune surveillance mechanism constantly in search of pathogens ^1^. Upon recognition of pathogens such as bacteria or viruses its activation results in a cascade of reactions ultimately leading to the lysis and killing of the identified pathogen^2^. Activation of the complement cascade may be regarded as a protective mechanism ultimately leading to the elimination of virus particles and disease progression. However, uncontrolled or prolonged complement activation may have a negative effect contributing to disease pathogenesis.

We have previously reported overactivation of complement in patients with severe and critical COVID-19 in line with similar studies which also showed elevated levels of anaphylatoxins C3a and C5a and increased circulating C5b-9 levels in serum and plasma patient samples ^3, 4, 5^. Furthermore, C3a and C5b-9 have been reported as independent predictors of COVID-19 severity and mortality^3,6^. Significant deposition of C4d, C5b–9 as well as mannan-binding lectin serine protease 2 (MASP2) has been detected in skin and lung samples of COVID-19 patients with a co-localisation of C4d and C5b-9 with SARS-COV-2 spike protein^7^.

Complement regulatory proteins (CRPs) CD55, CD59, CD46, and CR1 are all cell surface proteins which act to control complement cascade activation by cleaving complement factors and halting pathway activation (^8^). Specifically, CR1, CD55 and CD46 act at both the C3 and C5 steps promoting decay of C3 and C5 convertases whereas CD59 acts exclusively at the final step preventing formation of the membrane attack complex (MAC) complex and therefore cell lysis ^9^.

Taken together the above observations raise the question of whether a dysregulated expression pattern of CRPs may be present in COVID-19 contributing to complement overactivation. Using single-cell RNA sequencing (scRNA-seq) on PBMC isolated from COVID-19 patients of different disease states we characterised the immune response, assessed the expression signatures of CRPs in the various cellular populations and investigated the probable involvement of CRPs in COVID-19 pathogenesis.

## Results

### PBMC transcriptomic landscapes of COVID-19 patients

Single-cell transcriptomic profiles were generated from peripheral blood mononuclear cells (PBMCs) of COVID-19 patients with mild, severe and critical disease. Blood samples were obtained within 24 hours of admission and processed immediately. All critically ill patients were intubated. Healthy volunteers served as controls. Patients shared typical COVID-19 characteristics such as increased C-reactive protein and a marked lymphopenia (Figure 1b) in line with previous reports ^10,11^. Using scRNA-seq analysis we initially identified the PBMC transcriptomic profiles of healthy controls and COVID-19 patients. A total of 61,022 single cells passed quality control and were used in the analysis. The mild group consisted of 22,570 sequenced single cells while the severe and critical groups consisted of 7,350 and 15,397 cells respectively. Moreover 15,705 single cells were sequenced in the healthy group.

**Figure 1.**
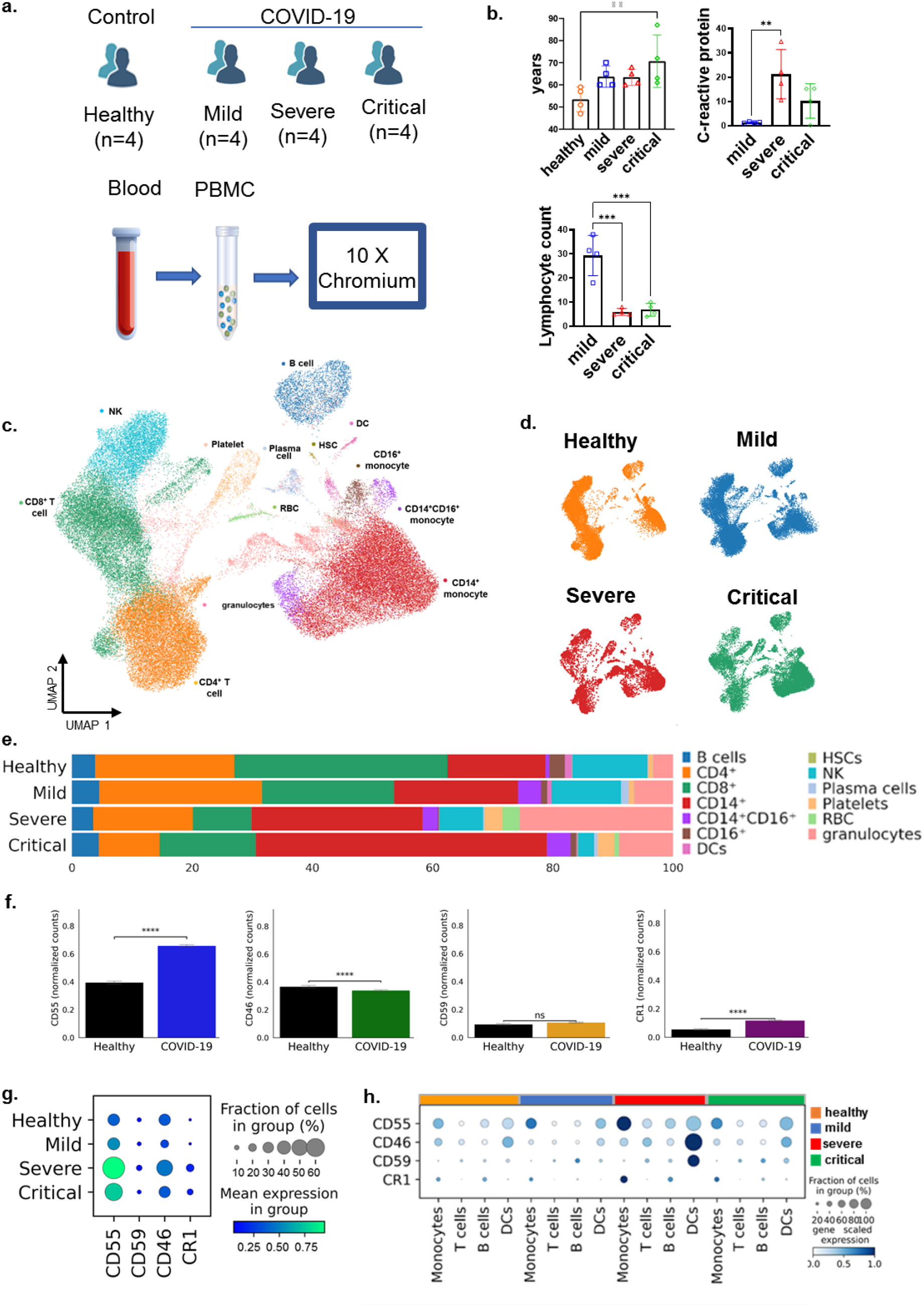
Single cell RNA sequencing analysis of PBMCs isolated from COVID-19 patients and healthy donors. a) Overview of the study participants and schematic representation of the experimental design for single cell RNA sequencing analysis of PBMCs isolated from blood samples of mild (n=4), severe (n=4), critical (n=4) COVID-19 patients and healthy controls (n=4). b) Demographic and clinical characteristics of patient groups and healthy controls. c) Major immune cell subsets identified by UMAP visualization, d) Transcriptomic profiles of healthy controls and mild, severe, critical COVID-19 patient PBMCs e) Bar plot of cell subsets presented in c and their changes according to disease state, expression levels of CRP proteins between healthy controls and COVID-19 patients (f), in the PBMC population of healthy and COVID-19 patients of different stages (g), in PBMC cells subsets (monocytes, T cells, B cells, DCs) according to disease state (h). Statistical analysis performed by the Wilcoxon rank sum test ****p<0.0001.

Integrated analysis of all the samples revealed 13 transcriptionally distinct PBMC populations (Figure 1c) and identified alterations between healthy controls and COVID-19 patients, as well as differences of cell states within each population (Figure 1d). PBMC populations were initially annotated manually based on the expression of standard markers used for PBMC subpopulation discriminatory analyses. Characterization of PBMC populations was performed according to the following markers: T-cells: CD3 (*CD3D, CD3G*) and *MAL* expression and *CD8A* for CD8^+^ cells, B-cells by *MS4A1* (CD20) and CD19 expression, natural killer cells (NK) by *KLRF1* and *NKG7* expression (Supplemental Figure 1). Monocytes were characterized by expression of *CD14* and *FCGR3A* (CD16^+^) expression, respectively, and erythrocytes by *HBA1* expression in combination with other genes. Validation of our annotation was performed against deposited datasets by COVID-19 patients PBMC analyses ^12,13^. Distinct locations for CD4^+^ and CD8^+^ T cells, natural killer (NK cells), CD14^+^ (classical), CD14^+^CD16^+^ (intermediate) and CD16^+^ (non-classical) monocytes, B-cells, dendritic cells (DCs) and plasmacytoid DCs (pDCs), granulocytes/neutrophils, platelets, hematopoietic stem cells (HSCs) and red blood cells (RBCs) were observed on the UMAP (Figure 1c). Integration analysis revealed an expansion of CD14^+^ monocytes, in severely and critically ill patients with a concurrent decrease of CD16^+^ (Figure 1d and 1e). A marked shrinkage was also observed in the T lymphocyte UMAP space in severe and critical patients, compared to the mild COVID-19 group of patients and healthy controls, (Figure 1d and 1e) attributed to both CD4^+^ and CD8^+^ T cells in line with the cytopenia observed in COVID-19 patients ^11^ as well as to NK cells (Figure 1d and 1e). B-cells remained unaltered quantitatively within disease states with only a mild reduction observed in severe COVID-19 patients (Figure 1d and 1e).

We initially investigated the expression levels of CRPs in the total PBMC populations of healthy and COVID-19 patients and observed an increase of CD55 (p<0.0001), CD59 (p=NS) and CR1 (p< 0.0001) expression in COVID-19 patients versus healthy controls and a slight decrease of CD46 expression (p<0.0001) (Figure 1f). Subsequently we determined the CRP expression in PBMCs of COVID-19 patients (Figure 1g). Similar expression levels of CD55 and CD46 were observed in PBMCs of healthy individuals while CD59 and CR1 were weakly expressed in comparison to CD55 and CD46. The relatively low expression of CD59 was also observed in whole blood samples of healthy individuals (Supplemental Figure 1). A trend of increased expression in severely ill COVID-19 patients followed by a relative decrease in critically ill was observed for all CRPs (Figure 1g) while the low expression levels observed for CD59 and CR1 in comparison to CD55 and CD46 were maintained in all disease states and controls (Figure 1f).

We next aimed at identifying the cell-type-dependent expression of membrane-bound complement regulators CD55, CD46, CD59 and CR1 in PBMCs and how this changed according to the disease state examined. To this end, CRP expression profiles were analysed in healthy controls as well as COVID-19 patients at different disease states (Figure 1h). As shown in Figure 1h, expression of CD55 and CD46 in healthy individuals was higher in comparison to CD59 and CR1 expression levels in all cell types. Expression of CRPs differed according to cell type in healthy controls with CD55 maximal expression in monocytes followed by DCs and B cells and lowest CD55 expression levels in T cells. CD46 was predominantly expressed in DCs followed by monocytes while similarly low levels of expression were observed in T cells and B cells. CD59 expression was minimal in all PBMC subpopulations compared to CD55 and CD46 with the lowest expression detected in monocytes while CR1 expression was higher in monocytes and B cells (Figure 1h) compared to T cells and DCs in which expression was barely detectable.

CRP expression varied according to COVID-19 state (Figure 1h). Specifically, an increase of CD55 expression was observed in monocytes and DCs in both mild and severe COVID-19 patients followed by a slight decrease of expression in monocytes of critically ill patients. The same trend was also observed in T and B lymphocyte subsets (Figure 1h). A slight reduction of CD46 expression levels was observed in mild COVID-19 patients, in all cell subsets, followed by an increase in severely ill patients especially in DCs. CD46 expression returned to levels similar to those of healthy individuals in critically ill COVID-19 patients (Figure 1h). CD59 expression remained at low levels in all cell types at all disease states apart from DCs of severely ill patients in which a striking increase was observed (Figure 1g). CR1 expression was also maintained at low levels in comparison to CD55 across all disease states and all cell types with a relative increase in monocytes of severely ill COVID-19 patients (Figure 1h).

### Transcriptome profile changes in the monocyte population of COVID-19 patients

The observed trend of increased CRP expression in severe COVID-19 patients followed by a decline in critically ill prompted us to investigate whether this was maintained within each cellular subpopulation or if it was dependent on specific cell traits. Therefore, we explored further the expression patterns of CRPs in specific cellular subpopulations by performing integrated subclustering analysis. Initial integrated subclustering analysis of all monocytes identified classical (CD14^+^), intermediate (CD14^+^CD16^+^) and non-classical (CD16^+^) monocytes and a total of 13 clusters (Figure 2a and 2b). A major expansion in the CD14^+^ super-cluster with a concurrent decrease of the CD16^+^ cluster was observed in the severe and critically ill states (Supplemental figure 2a). Specifically, an expansion of inflammatory monocytes CD14^+^IL1B (*CCL4, IL1B, CCL3*) and CD14^+^MPIG6B (*MPIG6B, GP9*) potentially affecting megakaryopoiesis was observed in critically ill patients (Figure 2c, 2d and 2f). Interestingly, cluster CD14^+^IL1B shared similarities with CD14^+^CD16^+^CXCL10 cluster (*TNF, CXCL10*,) of intermediate monocytes with a highly inflammatory profile which was only present in COVID-19 patients and decreased in critically ill patients (Figure 2c, 2d and 2f). Moreover, intermediate monocyte cluster CD14^+^CD16^+^RORA (*RORA, TXK, IL7R, BCL11B*) exhibiting macrophage like characteristics ^14^ diminished in severely and critically ill patients probably as a result of macrophage differentiation and tissue migration (Figure 2c, 2d and 2f). The third intermediate monocyte cluster CD14^+^CD16^+^ C1Q (*C1QC, C1QA, C1QB*) bared *C1QA* and *C1QB* common non-classical monocytes markers^15^. However it still exhibited C1QC, an intermediate monocyte marker, and expanded in critically ill individuals. CD14^+^LTF (*LCN2, MMP8, MMP9, LTF*) classical monocytes cluster was only observed in severe and critically ill COVID-19 patients and also displayed macrophage-like features (Figure 2c, 2d and 2f). The only non-classical cluster identified CD16^+^ (*LYPD2, CDKN1C, HES4, VMO1*) ^15^ progressively decreased in all groups of COVID-19, possibly due to their highly mature status and proapoptotic tendency. Two classical monocyte clusters CD14^+^MAP2K6 and CD14^+^SAP30 sharing a highly anti-inflammatory profile ^16^ also follow a similar trend of expansion in severe COVID-19 and further decline in critically ill patients (Figure 2c, 2d and 2f).

**Figure 2.**
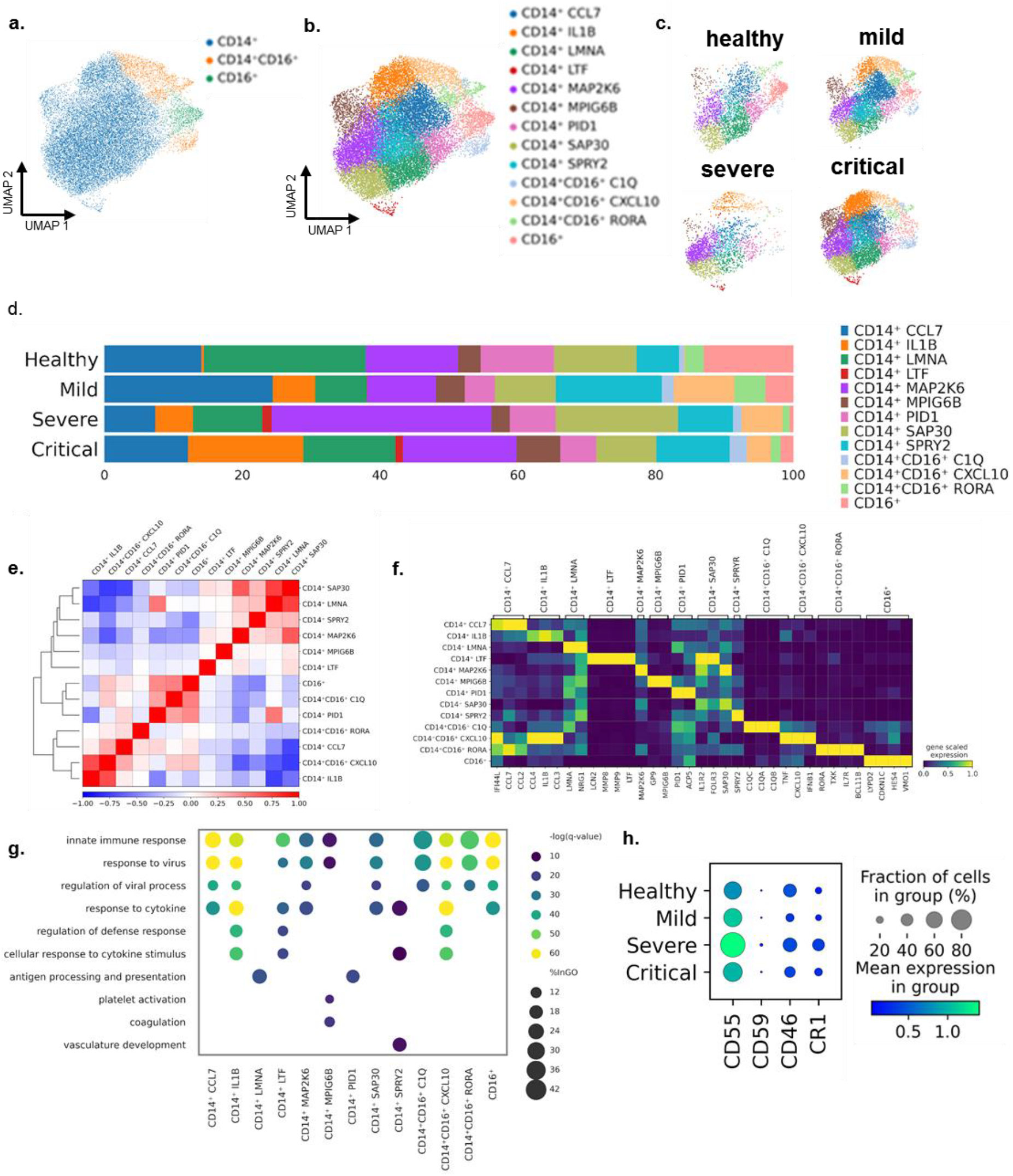
Compositional changes and expression of complement regulatory proteins in monocytes subpopulation. Integrated clustering analysis of 19.303 monocytes projected in the UMAP space a) clustering inferred from CD14^+^ and CD16^+^ expression b) identified monocyte subpopulations c) disease state specific monocyte subpopulations projected in the UMAP space (healthy group: 3,060 cells, mild disease group: 5,750 cells, severe disease group: 2,289 cells and critical disease group: 8,204 cells) d) Bar plot representing subpopulation quantitative changes (%) in each disease state, e) Correlation plot obtained from CD14^+^ monocyte subclustering analysis. f) Matrix plot of selected marker genes for each monocyte subpopulation characterisation and annotation g) Integrated functional analysis (GO, Coronascape) for each monocyte subpopulation h) Dot plot depicting expression levels of CRPs in all the monocyte subpopulations in each disease state.

Functional, gene ontology (GO), analysis revealed similar profiles related to innate immune responses, viral and cytokine responses and regulation of viral and defence responses shared between all clusters apart from CD14^+^LMNA and CD14^+^PD1 displaying a profile associated with antigen processing and presentation and cluster CD14^+^MPIG6B exhibiting a coagulation and platelet activation related profile (Figure 2g).

CRP expression was subsequently assessed in the monocyte states continuum. CRPs expression was detected in all monocyte clusters (Supplemental Figure 2b and 2c). However, in line with our observation in the global PBMC atlas (Figure 1f and 1g), CD59 expression was strikingly low in all COVID-19 severity states, suggesting a potential vulnerability to increased complement activation and possibly deposition due to the low levels of expression of the inhibitor of MAC formation (Supplemental Figure 2b and 2c). CD55 expression levels were relatively the highest, followed by CD46, and CR1. We then investigated CRP expression in COVID-19 disease states and observed an increases expression of CD55 and CR1 in all COVID-19 patients compared to healthy controls with the most profound increase in severe patients (Figure 2h). On the contrary, CD46 expression levels remained similar between severe COVID-19 patients and healthy controls. A slight decrease in expression of all CRPs was observed in critical patients compared to severe (Figure 2h) in line with the trend observed in whole PBMCs (Figure 1g). The expression pattern observed for CD55 in our data was further confirmed by verification in a large publicly available dataset of scRNA-seq data^12^. The dataset was generated by analysis at the single cell level of PBMCs isolated from a total of 102 COVID-19 patients of various states. A similar trend of increased expression in severe COVID-19 followed by a decline in critical COVID-19 patients was also observed in the large dataset for CD55 and CR1 thus confirming our observations, while the marginal level of CD59 expression did not allow for direct comparison (Supplemental figure 3). Finally, we confirmed the decrease in CD46 expression observed in critical COVID-19 patients in comparison to healthy.

**Figure 3.**
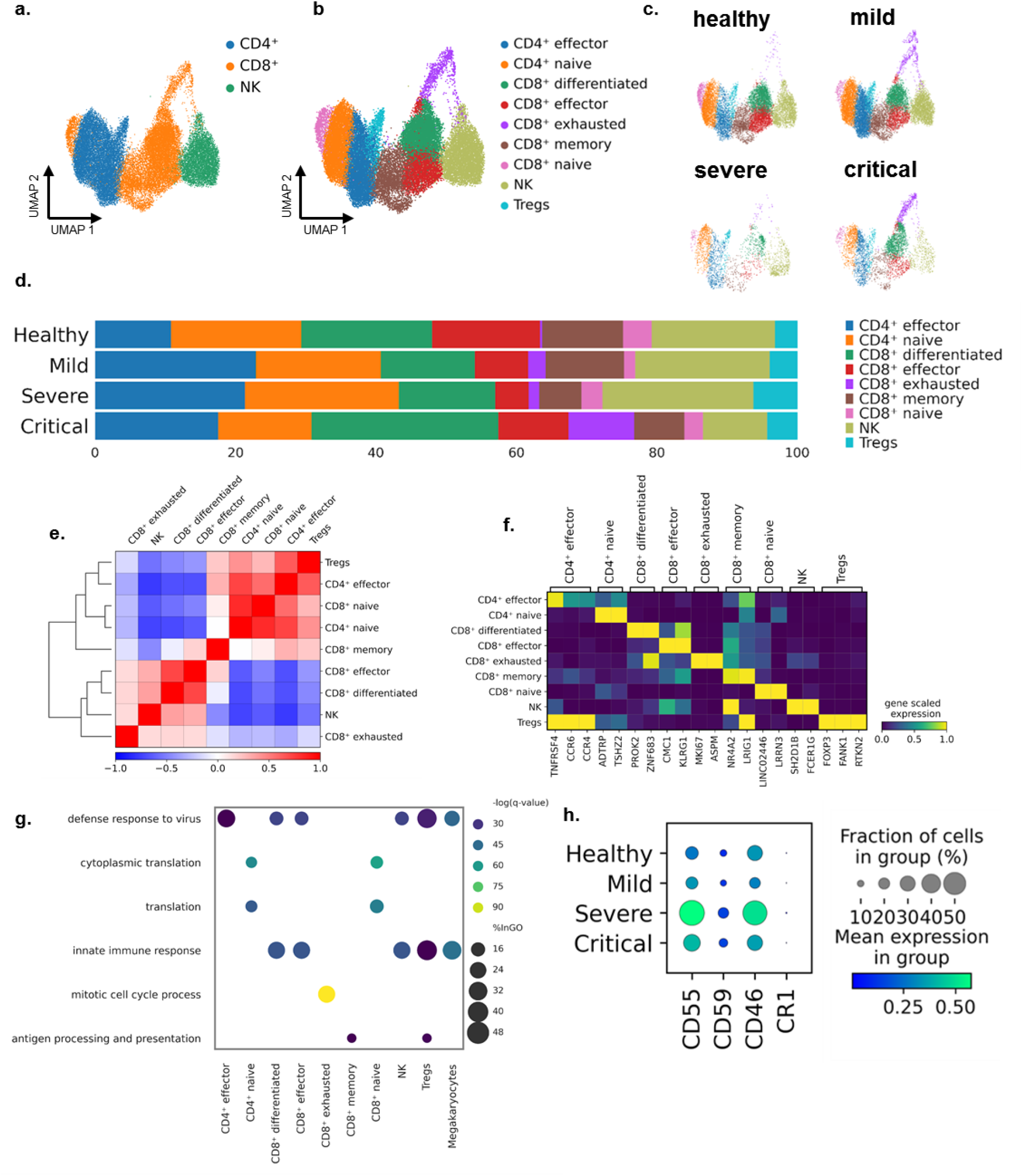
Compositional changes in T cells subpopulations of COVID-19 patients. Integrated clustering analysis of 31,763 T cells projected in the UMAP space a) clustering inferred by CD4^+^, CD8^+^ and NK expression marker genes b) identified T cell subpopulations c) disease state specific T cell subpopulations projected in the UMAP space (healthy group: 11,180 cells, mild disease group: 13,687 cells, severe disease group: 2,470 cells and critical disease group: 4,426 cells) d) Bar plot representing subpopulation quantitative changes (%) in each disease state, e) Correlation plot obtained from the integrated T cell subclustering analysis. f) Matrix plot of selected marker genes for each monocyte subpopulation characterisation and annotation g) Integrated functional analysis (GO, Coronascape) for each T cell subpopulation h) Dot plot depicting expression levels of CRPs in all the T cell subpopulations in each disease state.

We next analysed the observed decrease of CD55 expression. CD55 displayed a statistically significant decrease of expression in classical CD14^+^ monocyte subpopulations (Supplemental Figure 2d) of critical COVID-19 patients compared to severe, suggesting an unsustained inhibition of complement overactivation. However, CD16^+^ monocytes tend to maintain expression of CRPs in order to achieve inhibition of complement overactivation. Interestingly, decreased CD55 expression may be attributed to distinct subpopulations. Specifically, a statistically significant reduction of CD55 was only observed in all CD14^+^ subpopulations, which shared a highly inflammatory profile at the functional level (Supplemental Figure 2e).

### Transcriptomic alterations in T-cell populations of COVID-19 patients

Integrated analysis of the T cell subpopulation generated 10 distinct supopulations including CD4^+^ and CD8^+^ naïve and inflammatory cells, cytotoxic effector cells, Tregs, NK and proliferative CD8^+^ T cells (Figure 3a and b). Specifically, an expansion of effector CD4^+^ T cells with a highly inflammatory profile (*TNSF4, CCR4, CCR6*), as well as in differentiated CD8^+^ T cells responsive to viral infection and IFN-γ (*PROK2, ZNF683*) was observed in all COVID-19 disease states compared to healthy controls (Figure 3b, 3d and 3f) (Figure 3b, 3d and 3f) ^17 18^. A cluster of highly proliferative cytotoxic CD8^+^ T cells (exhausted CD8^+^ T cells) was mostly observed in COVID-19 patients, expanded in critically ill, and bared markers indicative of exhaustion (*MKI67, ASPM*) ^19^ (Figure 3b, 3d and 3f). An expansion of T-regs (*FOXP3, FANK1, RTKN2*) ^19^ in severe COVID-19 patients was observed with a subsequent reduction in critically ill (Figure 3b, 3d and 3f). CD8^+^ effector cells (*CMC1, KLRG1*) ^20^ were shown to decrease in severe and critically ill patients possibly as a result of differentiation and *KLRG1* loss. A similar decrease was observed for another cluster of memory CD8^+^ T cells (*NR4A2, LRIG1*) with exhaustion features (*GZMK, CCL5*) ^21^ (Figure 3b, 3c and 3f).

GO analysis revealed enriched functions related to defence responses to viral infections and innate immune responses for almost all T cell subpopulations, except for naive CD4^+^ and CD8^+^ subpopulations which were characterised by a profile associated with translation. Exhausted CD8^+^ T cells bared a profile of mitotic cell cycle processes while the CD8^+^ memory subpopulation exhibited a profile related to antigen processing and presentation (Figure 3g).

We next assessed the expression of CRPs in all T cell sub-populations (Figure 3h). As shown in Figure 3h CD55 and CD46 exhibited relatively the highest expression levels while minimal expression was observed for CD59 and CR1 expression was practically absent. The CRP expression profiles in the T cell subpopulation detected in our dataset were also validated in the larger published datasets ^12^ (Supplemental Figure 3b). In line with our observations, CD55 upregulation was confirmed in the severe group of patients of the larger dataset (Supplemental Figure 3). However, in contrast to the decrease of CD55 expression observed in the critically ill patient group of our dataset, a further increase of CD55 was shown in the larger dataset. Finally, the expression of CD46 and CD59 follow the same trend in both datasets examined.

### Transcriptomic alterations in B-cell populations of COVID-19 patients

Subclustering analysis of the B cell population resulted in its discrimination into seven distinct subopulations including B cells (*CRIP1, CIB1*), naïve B cells (*TCL1A*) ^22^, two mature B cell subpopulations (*GPR183, MARCS, TBC1D9, LNC01857*) ^23 24^, memory B cells (*YBX3, AFF3, FCER2*) ^25^, plasma cells (*JCHAIN, TXNDC5*) and plasmablasts (*RRM2, MKI67*) (Figure 4c and 4e). Mature B cells showed a reduction in severe COVID-19 patients with a direct expansion in critically ill patients while the opposite was observed for memory B cells which showed a massive expansion in severe COVID-19 patients and a decrease in critically ill (Figure 4c and 4e). Naïve B cells and plasma cells remained fairly unaltered within each disease state while plasmablasts were only observed in COVID-19 patient profiles specifically in the mild and critical COVID-19 groups of patients (Figure 4c and 4e) possibly as the sources of antibodies production.

**Figure 4.**
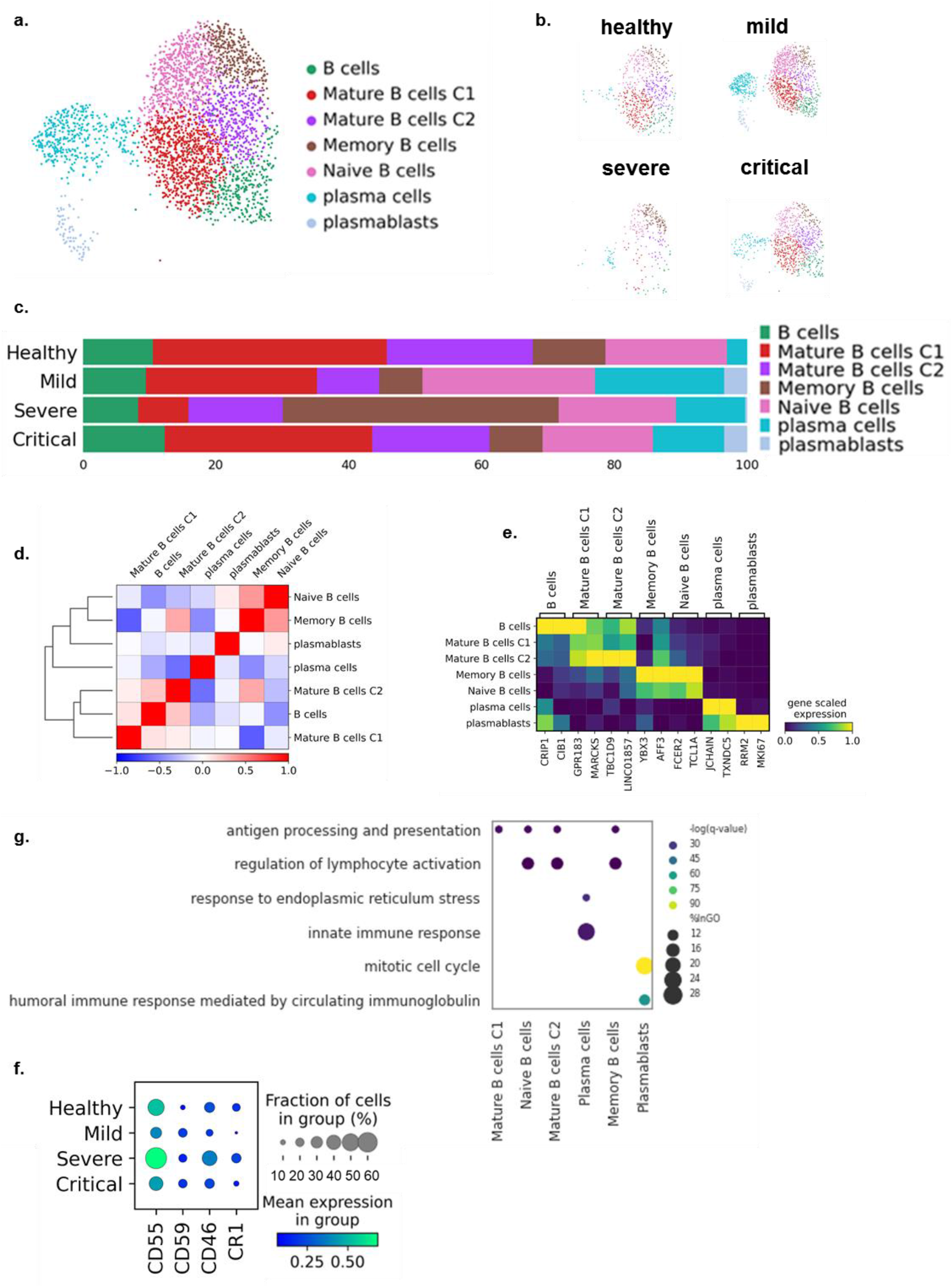
Compositional changes in B cells subpopulation of COVID-19 patients. Integrated clustering analysis of 3,038 B cells projected in the UMAP space a) Identified B cell subpopulations b) disease state specific B cell subpopulations projected in the UMAP space (healthy group: 617 cells, mild disease group: 1,328 cells, severe disease group: 289 cells and critical disease group: 804 cells) c) Bar plot representing subpopulation quantitative changes (%) in each disease state, d) Correlation plot obtained from the integrated B cell subclustering analysis. e) Matrix plot of selected marker genes for each monocyte subpopulation characterisation and annotation g) Integrated functional analysis (GO, Coronascape) for each B cell subpopulation h) Dot plot depicting expression levels of CRPs in all the B cell subpopulations in each disease state.

GO analysis in the B cells population revealed profiles related to antigen processing and presentation as well as to regulation of lymphocyte activation for the naïve and mature B cells. Functions related to innate and humoral immune responses were identified for the plasma cells and plasmablasts respectively (Figure 4g).

Analysis of the expression of CRPs in the B cells population revealed relatively higher CD55 expression levels in healthy individuals in comparison with other CRPs (Figure 4f). Lower CD55 expression levels were observed in the mild COVID-19 group of patients with a marked upregulation in the severe COVID-19 patients. Finally, CD55 expression levels returned to levels similar to those of the mild COVID-19 group in critically ill patients (Figure 4f). CD46 and CR1 expression followed the same trend and CD59 showed an upregulation that remained fairly stable in all COVID-19 stages examined (Figure 4f). Verification of the CRPs expression pattern in the larger published dataset, revealed an unexpected decrease in the severe COVID-19 group (Supplemental figure 3b). However, both CD46 and CD59 expression levels were confirmed while CR1 expression was too weak to assess quantifiable results (Supplemental figure 3b).

### Expression patterns of CD55 and CD59 proteins in monocytes, T cells and B cells of COVID-19 patients by fluorescence activated cell sorting (FACS) analysis

The transcriptomic profiles observed for CRPs expression in COVID-19 patients revealed an increase of CRPs in severely and critically ill patients compared to controls with a trend of a reduced expression in the critical COVID-19 group compared to severe COVID-19 patients in both monocytes, T cells and B cells (Figures 2h, 3g and 4f). In order to assess whether the transcriptomic profiles obtained for CRPs in COVID-19 could be verified at the protein level FACS analysis for the expression of CD55 and CD59 was performed. We assessed their expression in monocytes, T cells, B cells and granulocytes (neutrophils and eosinophils) separately. We focused primarily on CD55 and CD59 as the former CRP acts early on in the cascade and CD59 acts solely at the final step of the cascade thereby limiting C5b-9 or membrane attack complex (MAC) formation and deposition and therefore cell lysis. A statistically significant increase of CD55 expression (p<0.05) in the severe COVID-19 group of patients (p<0.05) following a significant decrease in critically ill was observed in monocytes in line with the trend observed in our single-cell transcriptomic analysis (Figure 5a). A significant increase in CD55 expression was observed in CD4^+^ T cells in severely (p< 0.01) and critically ill (p<0.001) COVID-19 patients compared to healthy controls. CD8^+^ T cells, showed significantly increased CD55 expression in the severe versus healthy (p<0.05), critical versus healthy (p<0.0001) and critical versus severe (p<0.01) COVID-19 groups of patients (Figure 5a). This increase in CD55 protein expression levels is in line with our single-cell transcriptomic data analysis (Figure 3). B cell CD55 expression was significantly increased in the critical COVID-19 group of patients compared to severely ill (p<0.05) and healthy controls (p<0.05). No significant changes were observed in granulocyte (neutrophil and eosinophil) CD55 expression. A significant increase in CD59 monocyte and granulocyte expression was observed in severely and critically ill COVID-19 patients compared to healthy controls with no significant change in CD4^+^ and CD8^+^ T cell CD59 expression. CD59 was significantly reduced in B cells of critical COVID-19 patients compared to the severe COVID-19 group (Figure 5b).

**Figure 5.**
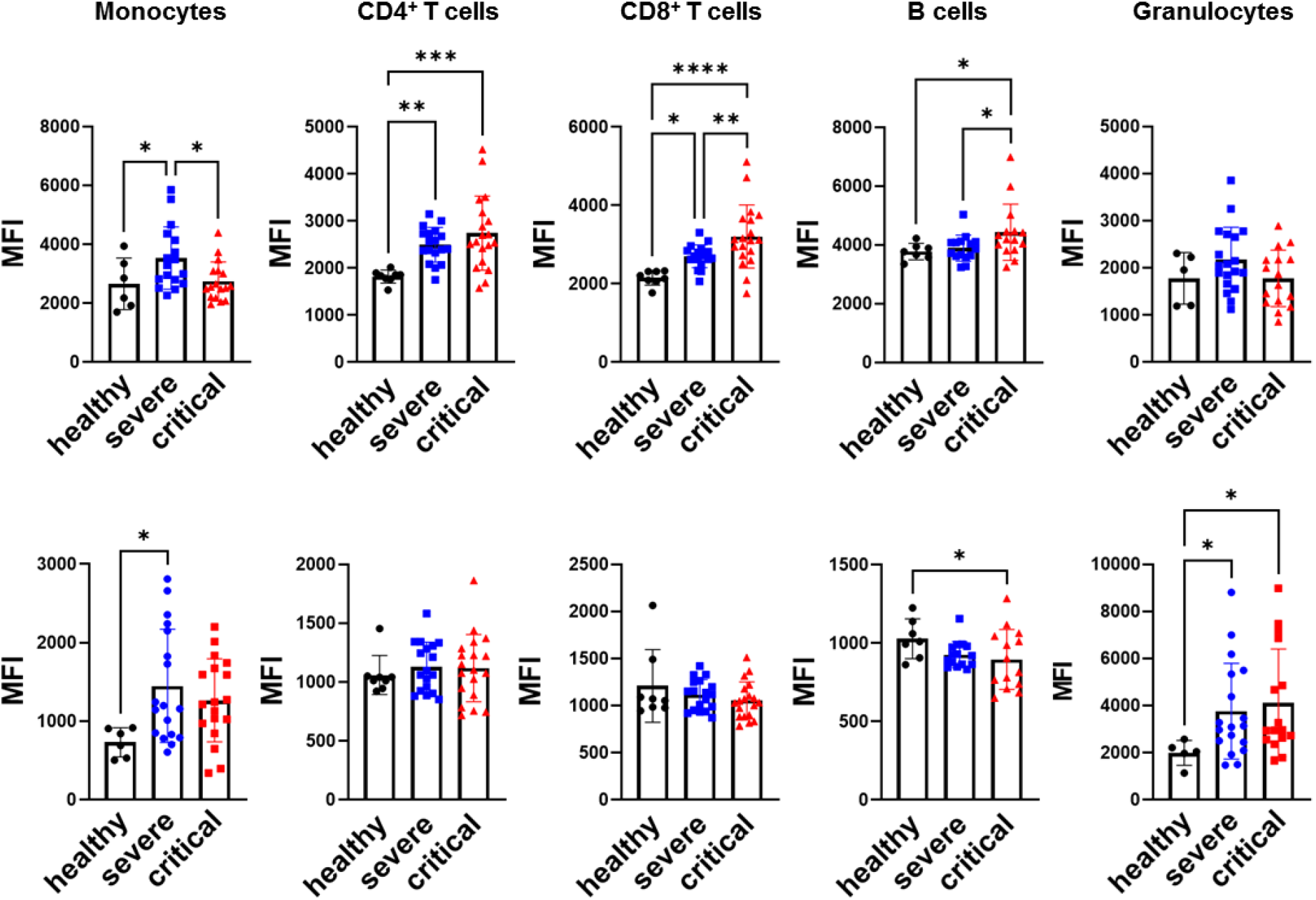
Validation of CRP expression patterns by flow cytometric analysis. a) Flow cytometric (FACS) analysis was carried out in whole blood samples of COVID-19 patients with severe (n=20) and critical (n=20) disease in order to determine expression levels of CD55 (a) and CD59 (b) in monocytes, CD4^+^ T cells, CD8^+^ T cells, B cells and granulocytes Statistical analysis was performed by One way ANOVA testing for more than two group comparisons and by the Kruskal-Wallis non-parametric test when normality of data was not achieved. Post-hoc analysis was performed by the Least significant difference (LSD) test. *p<0.05, **p<0.01, ***p<0.001, ****p<0.0001.

### CD55 upregulation in severe and critical COVID-19 patients corresponds with suppression of IFN responses

The upregulation of CD55 observed in the severe group of COVID-19 was further confirmed by bulk sequencing analysis of PBMCs (Supplemental figure 4a) and by Real-time PCR amplification on whole blood samples albeit the latter was not significant possibly due to the presence of other blood cells which may mask the increase of CD55 expression observed in PBMC (Supplemental figure 4b). Receiver operating characteristic analysis for the association of CD55 with prediction of disease progression such as admission to ICU revealed a high AUC value (AUC= 0753, p=0.002) supporting a probable important role of CD55 in disease pathogenesis (Supplemental figure 4c).

A previous study on profiling of CRP expression as potential markers of viral infection, other than SARS-COV-2, reported no change of CD55 expression in monocytes, lymphocytes or neutrophils with an increase of CD59 expression in monocytes and neutrophils compared to healthy controls, and unaltered CD59 expression levels in lymphocytes of virus infected patients compared to healthy controls ^26^. Therefore, the increase of CD55 expression observed, in our study, in the severe COVID-19 groups of patients by both scRNA-seq and FACS in monocytes and T cells suggests a distinct COVID-19 specific, CD55 expression. This distinctive expression pattern of CD55 observed in severe and critical COVID-19 patients prompted us to investigate whether it may exert an effect on immune responses of COVID-19 patients. Apart from its role in controlling the activation of the complement cascade, CD55 has been reported to possess various roles as an immunoregulator. Specifically, CD55 has been previously identified as a suppressor of T cells immune responses ^27^. We therefore questioned whether an effect of suppressed IFN responses could be identified in PBMC cell types at the stages of COVID-19 in which we observed CD55 overexpression. In order to achieve this, the expression of interferon stimulated genes (*IFI6, IFI27, IFITM3, IFI44L, IFIT1, IFIT3, STAT2, ISG15* and *ISG20*) (ISGs) was assessed in monocytes, T cells and B cells. The selection of the specific genes was based on the lists of genes obtained following differential expression analysis between COVID-19 states for each individual cluster and further confirmed by recent related studies ^28 13 29^. As shown in Figures 6a and 6b we observed an increase of ISGs expression in monocytes of the mild COVID-19 group followed by a sharp decline in both the severe and critical COVID-19 groups. The decline in ISGs expression coincided with the increase of CD55 expression suggesting a possible contribution to the suppression of IFN responses previously reported for these patients ^29^. Analysis of CD55 expression levels in CD14^+^ and CD16^+^ monocytes revealed higher expression levels in CD14^+^ monocytes compared to CD16^+^ indicating that the suppression of ISGs could possibly be attributed mainly to CD14^+^ monocytes (Figure 6c). As shown in Figure 6d all classical CD14^+^ monocyte clusters showed the highest CD55 expression levels in the severe COVID-19 group of patients which coincided with almost absent ISG expression following the increase observed at the mild COVID-19 stage. The increase of CD55 expression in CD14^+^ clusters was statistically significant for all CD14^+^ clusters apart from CD14^+^IL1B (Figure 6e). Furthermore, the only non-classical CD16^+^ cluster identified also exhibited a significantly increased expression of CD55 in the severe state (Figure 6e) coinciding with almost complete absence of the ISGs expression. This was in striking difference to the mild stage for CD16^+^ monocytes in which ISGs were highly expressed with a weaker CD55 expression suggesting that CD55 also exerts the same effect in CD16^+^ monocytes (Figure 6d). Detection of the ISGs expression on the UMAP of the monocyte subpopulation showed the increased expression of ISGs in the mild COVID-19 patients while ISGs expression was only observed in a region of intermediate CD14^+^ CD16^+^ monocytes in which CD55 expression was relatively low in comparison to other areas on the UMAP (Supplemental figure 5a and b).

**Figure 6.**
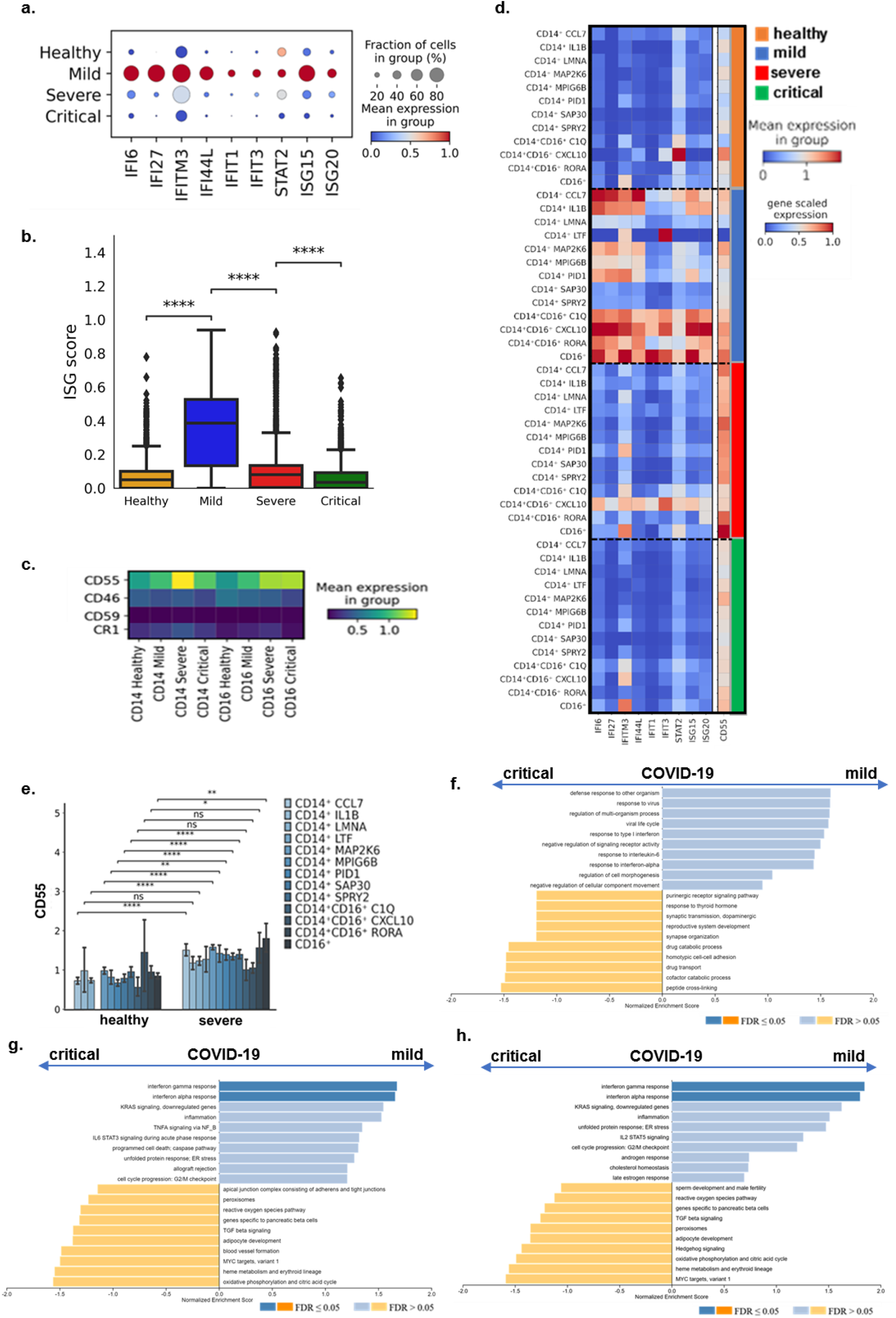
IFN responses are suppressed upon CD55 upregulation in monocytes of severe and critical COVID-19 patients. a) Dot plot depicting the expression levels of the ISGs in healthy controls and patients with mild, severe and critical COVID-19 b) box plot of ISG score changes in COVID-19 patients of different disease severity and healthy controls (Mann-Whitney-Wilcoxon test two sided with Bonferroni correction ****p<0.0001) c) Matrix plot representing the expression of ISGs and complement regulator CD55 in each monocytes subtype in COVID-19 patients, d) Matrix plot representing the expression of CRPs in CD14^+^ and CD16^+^ monocytes of COVID-19 patients e) CD55 expression levels in monocyte subpopulations of severe COVID-19 patients compared to healthy controls. Statistical analysis performed by the Wilcoxon rank sum test *p<0.05, **p<0.01, ****p<0.0001. Quantitative bar plots produced by gene set enrichment analysis (GSEA) performed on the gene lists derived by comparison of severe versus mild COVID-19 in the f) classical monocyte CD14^+^IL1B cluster g) intermediate monocyte CD14^+^CD16^+^CXCL10 cluster and h) non-classical monocyte CD16^+^ subpopulations in mild versus critical COVID-19 patients. FDR ≤0.05.

The decrease of IFN responses in severe patients was further confirmed by gene set enrichment (GSEA) analysis for specific clusters of classical monocytes, CD14^+^IL1B cluster, intermediate monocytes CD14^+^CD16^+^CXCL10 cluster and non-classical monocytes CD16^+^ cluster. A suppression of interferon responses was observed in critical COVID-19 patients compared to the mild group in which IFN responses were increased (Figure 6f, g, h).

We next examined whether a similar association of increased CD55 expression with suppressed interferon responses could be observed in the T cell population. Again, a major increase of ISGs was identified in the mild COVID-19 group of patients with a subsequent reduction in the severe and critical COVID-19 groups (Figure 7a and 7b). As shown in the dot blot of the ISGs expression, *IFI6, ISG15* and *ISG20* expression levels were the highest compared to the other genes in all groups of patients as well as in healthy controls and maintained in the severe COVID-19 group of patients. This was in contrast to the ISG expression pattern observed in monocytes in which expression levels of all ISGs were similar and followed the same expression pattern (Figure 6a). As shown in Figure 7b, ISG score expression levels in T cells gradually returned to those observed in healthy controls in the critical COVID-19 group of patients.

**Figure 7.**
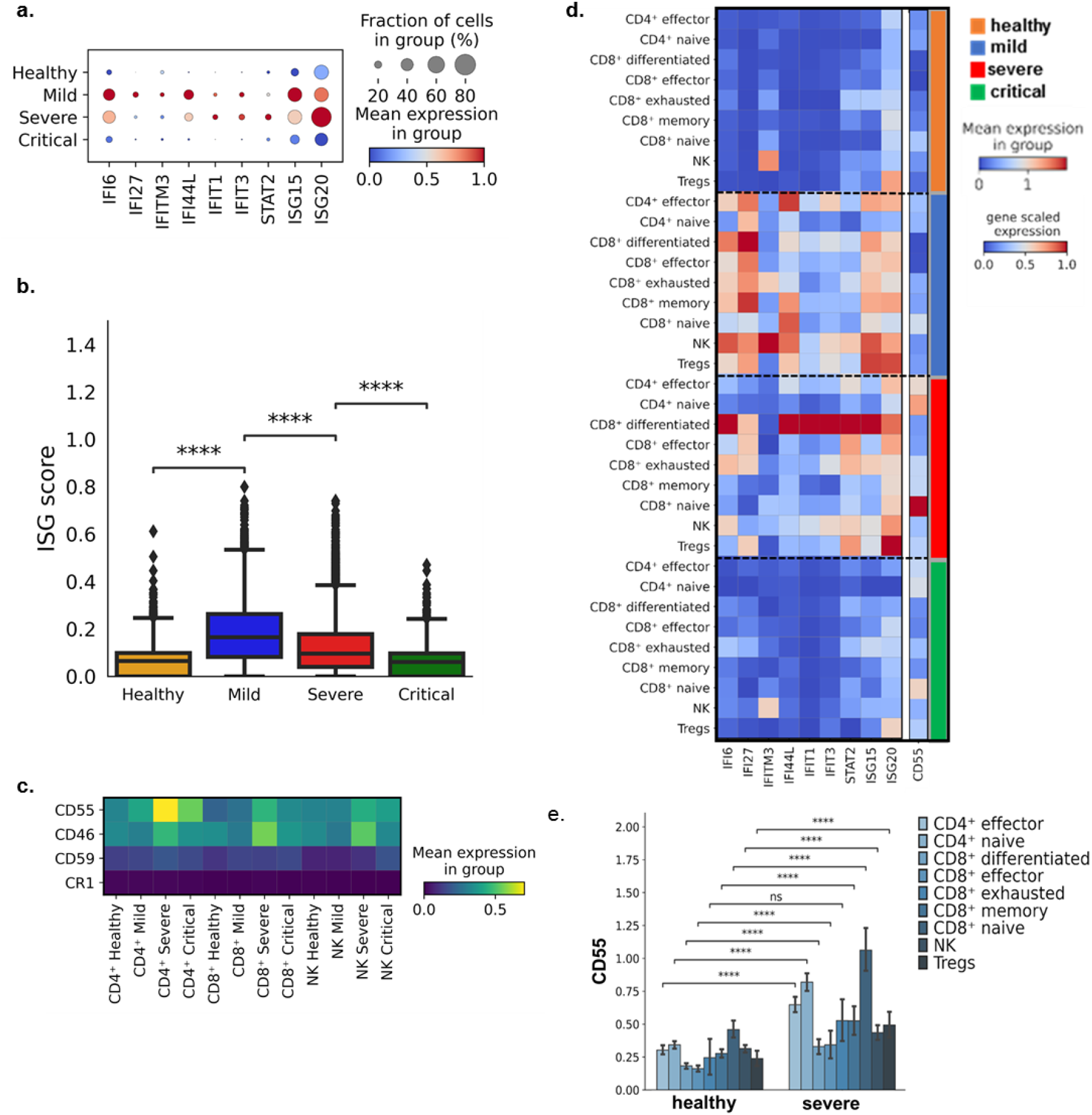
IFN responses are suppressed upon CD55 upregulation in T cells of severe and critical COVID-19 patients. a) Dot plot depicting the expression levels of the ISG score in healthy controls and patients with mild, severe and critical COVID-19 b) Box plot representing ISGs expression all the T cell subpopulations in all the COVID-19 disease state and healthy individuals (Mann-Whitney-Wilcoxon test two sided with Bonferroni correction, ****p<0.0001) c) Matrix plot showing CRPs expression in CD4^+^, CD8^+^ T and NK cells of COVID-19 patients d) Matrix plot representing ISGs and CD55 gene expression in individual T cell subpopulations of COVID-19 patients, e) CD55 expression levels in T cell subpopulations of severe COVID-19 patients versus healthy controls. Statistical analysis performed by the Wilcoxon rank sum test ****p<0.0001.

Analysis of expression of CD55 in CD4^+^ and CD8^+^ T cells and NK cells revealed a higher induction of CD55 expression in CD4^+^ T cells compared to CD8^+^ and NK (Figure 7c) suggesting that the reduction of the ISG score could be linked to the up-regulated expression of CD55 in CD4^+^ T lymphocytes. Indeed, in the severe COVID-19 group of patients CD4^+^ effector and CD4^+^ naïve clusters showed higher levels of CD55 expression in association with lower expression for almost all interferon response genes (Figure 7d) when compared to the mild COVID-19 group of patients. On the contrary, CD8^+^ differentiated, CD8^+^ effector and CD8^+^ memory clusters all exhibited lower levels of CD55 compared to the CD4^+^ effector and CD4^+^ naïve clusters and higher expression in the ISGs. Similarly, for the NK cell cluster which exhibited lower levels of CD55 in comparison to CD4^+^ clusters a higher expression for most ISGs was observed. The increase of CD55 expression in the severe COVID-19 group T cells subpopulation clusters was statistically significant for most clusters apart from the CD8^+^ exhausted cluster of cells (Figure 7e).

Finally, an association of reduced interferon response with increased CD55 expression was also identified in the B cell population. As shown in Figures 8a and 8b, the same trend of reduced expression of the ISGs in the severe and critical COVID-19 group of patients following an initial increase in the mild COVID-19 group was observed. Interestingly, ISG signatures were very similar to those observed in T cells with the highest expression levels observed again for *ISG15* and *ISG20* genes (Figure 8a) and the sustained expression of *ISG20* in the severe COVID-19 group probably due to their shared lymphocyte lineage. As shown in Figure 8c a direct decrease of ISGs expression could be associated with the increased CD55 expression for all B cell clusters in the severe group of COVID-19 patients. This follows the contrast in the mild group in which lower levels of CD55 allowed for increased ISG expression (Figure 8c). Expression levels of six (*IFI6, IFI27, IFITM3, IFI44L, IFIT1, IFIT3*) out of the nine ISGs returned to those observed in the healthy individuals and were almost undetectable. In contrast, *STAT2, ISG15* and *ISG20* displayed sustained higher expression levels in all clusters even in the critical COVID-19 groups of patients. Plasma cells and plasmablasts exhibited the highest expression levels for *ISG15* and *ISG20* in marked contrast to the other cell clusters (Figure 8d). The increase in CD55 expression was obvious in all clusters of the severe COVID-19 group of patients compared to healthy controls albeit it did not reach statistical significance.

**Figure 8.**
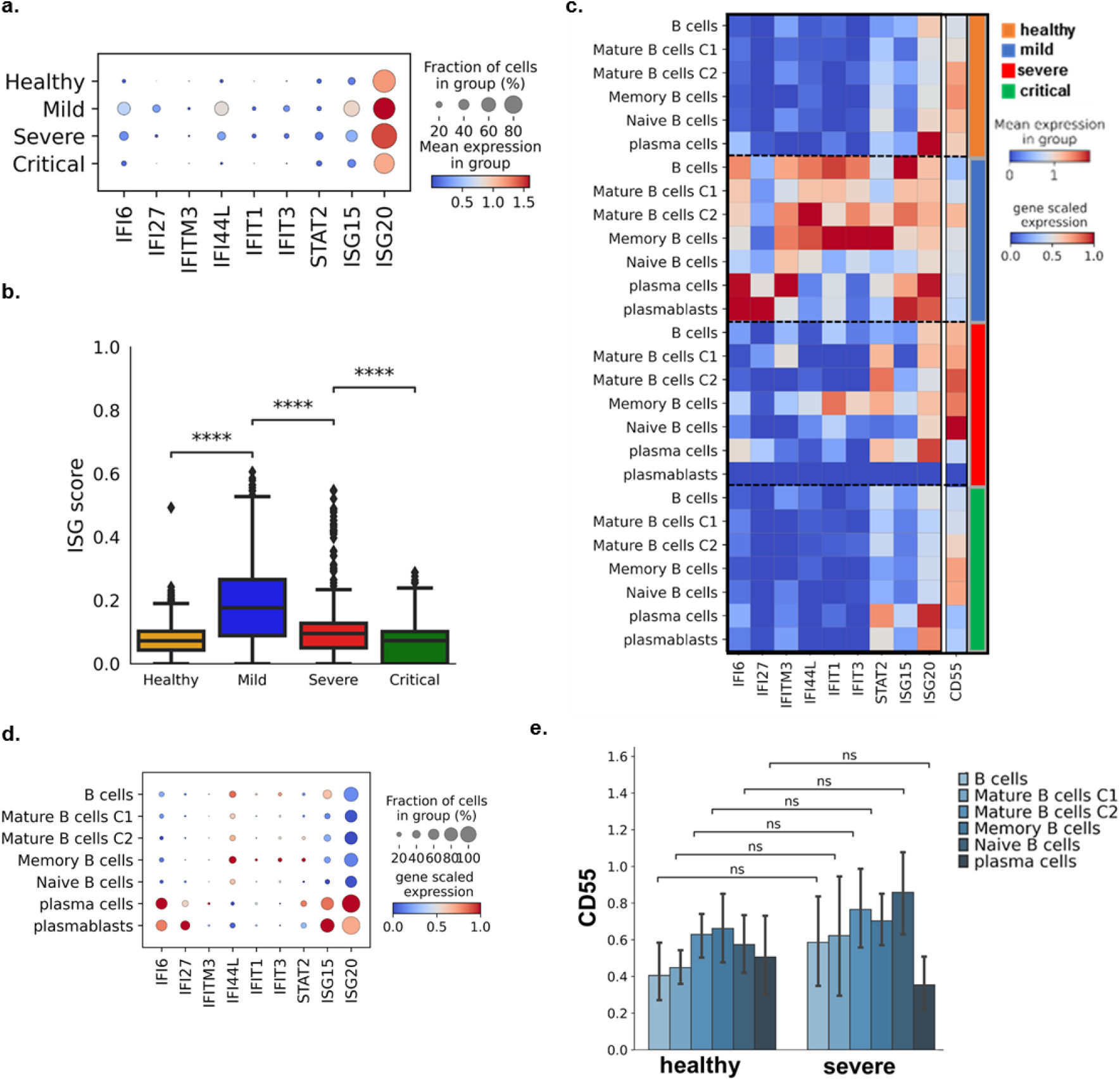
IFN responses are suppressed upon CD55 upregulation in B cells of severe and critical COVID-19 patients. a) Dot plot depicting expression levels of the ISGs in all the B cell subpopulations in healthy controls and patients with mild, severe and critical COVID-19 b) Box plot representing ISGs expression all the B cell subpopulations in all the COVID-19 disease state and healthy individuals (Mann-Whitney-Wilcoxon test two sided with Bonferroni correction, ****p<0.0001) c) Matrix plot representing ISGs and CD55 gene expression in individual B cell subpopulations of COVID-19 patients d) Dot plot depicting gene expression of ISGs indicated in each B cell subpopulation e) CD55 levels in B cell subpopulations of severe COVID-19 patients versus healthy controls. Statistical analysis performed by the Wilcoxon rank sum test, NS; non-significant.

## Discussion

Little is known about the role of CRPs during viral infections both through their regulatory activity on complement cascade activation and their presumed direct effect on antiviral processes and the immune response. Our sc-RNAseq study of PBMCs isolated from COVID-19 patients identified a distinct COVID-19 specific expression pattern for CD55 in severely and critically ill patients. A previous study which investigated the expression patterns of CD55, CD46, CD59 and CR1 in PBMCs in viral and bacterial infection versus healthy controls reported no effect on the expression levels of CD55 in monocytes, lymphocytes and neutrophils of patients with viral infection compared to healthy controls^26^. The study included 26 patients with a confirmed viral infection from various unrelated viruses including, Influenza A virus, Dengue virus, Hepatitis B virus, Epstein-Bar virus, Enterovirus, Herpes-simplex virus, Tick-borne encephalitis virus and others. In contrast, our scRNA-seq analysis revealed that PBMCs isolated from hospitalised COVID-19 patients, both severely and critically ill, displayed increased CD55 expression (Figure 1f and 1g). Up-regulated CD55 expression was further analysed in the individual subpopulations that we resolved by sc-RNAseq (Figure 2h, Figure 3g and Figure 4f). The upregulation of CD55 expression observed could be explained as a putative protective mechanism of blood cell subpopulations from uncontrolled complement activation. Increased levels of complement factors in COVID-19 patients have been reported by previous studies. De Noorja et.al reported increased levels of C3a, C3c and C5b-9 in COVID-19 patients in comparison to healthy controls ^4^ while Zolteck et.al reported significant elevation of C3a and C5b-9 in both hospitalised and critically ill patients compared to healthy controls as well as in non-survivors ^6^. Significantly high levels of circulating C5a and C5b-9 have also been reported in COVID-19 in comparison to influenza patients and ICU patients with non-COVID-19 respiratory failure ^30^, while others have shown increased deposition of complement factors in tissue samples from COVID-19 patients including lungs, liver, kidneys and the endothelium^7, 31^ thus rendering complement activation a distinctive feature of COVID-19. A recent study further demonstrated increased complement deposition in monocytes of COVID-19 patients of various disease states. Lange et.al identified significant elevation of C1Q and C3 deposition in the monocyte population of patients with severe and critical COVID-19 compared to healthy controls^32^. The upregulation of CD55 may therefore act to eliminate further activation and deposition of complement in this cellular population. In line with our observations, a recent study characterising CRP expression levels in monocytes of COVID-19 patients by FACS analysis ^32^ also reported an increase of CD55 levels in the severe COVID-19 group.

Furthermore, we observed an increase in the expression of CD59, which acts on the final step of the cascade inhibiting C5b-9 formation and deposition and therefore complement mediated cell lysis, in monocytes and granulocytes of patients with severe and critical COVID-19 with no upregulation in T cells or B lymphocytes. This expression profile was in accordance with the profile observed during viral infection ^26^. Therefore, the increase of CD55 observed in the specific cellular populations of COVID-19 patients with severe and critical disease may be taking place as an additional complement activation halting mechanism in order to achieve protection of these cell populations. Finally, we observed an upregulation of CD46 in monocytes as well as in T and B lymphocytes by scRNA-seq analysis. Augmented CD46 expression levels in monocytes have been reported in patients with viral infections with a concomitant downregulation of CD46 expression in T cells ^26^. Although not verified at the protein level the T cell CD46 upregulation may also present a COVID-19 specific expression for CD46. Taken together, the above observations indicate that the enhanced CD55 and CD46 expression levels observed in PBMCs during COVID-19 may signify an effort to activate the halting machinery of complement activation in order to protect PBMCs which are the first line of defence during the infection.

Interestingly, we also observed a reduction of CD55 expression in the monocyte population of the critical COVID-19 group of patients in comparison to the severe group (Figure 2h). This decrease was further confirmed by scRNA-seq in the dataset by Stephenson et.al (Supplemental Figure 3a) as well as by FACS analysis (Figure 5a) and may contribute to the sustained complement deposition reported in the monocyte population of convalescent individuals (Lange et.al).

The distinct feature of CD55 upregulation confirmed in monocytes and T and B lymphocytes of COVID-19 severe and critical patients was associated with reduced type I interferon responses (Figures 6, 7 and 8). Type I interferon responses are key mediators of the antiviral defence and involve IFN-a and IFN-β^33^. A suppression of type I interferon responses has been well documented in patients with severe and critical COVID-19. Hadjaj et.al demonstrated decreased expression of ISGs in patients with severe and critical COVID-19 following a marked increase in patients with mild disease^29^. Another study confirmed reduced ISG expression in monocytes and T lymphocytes by scRNA-seq in patients with mild COVID-19 compared to severely ill^28^. Wilks et.al further demonstrated reduced expression of type I interferon related genes in critical COVID-19 patients by scRNA-seq in monocytes and T cells^13^. The study revealed a huge variation in the profiles of type I interferon related genes in critical COVID-19 patients ranging from total absence of expression to upregulation^13^. We also confirmed the same profile of reduced type I interferon responses in the severe and critical COVID-19 groups of patients following a marked increase in the mild COVID-19 group by analysis of the expression of nine ISGs in both total PBMCs as well as in the monocyte, and T and B lymphocyte subpopulations (Supplemental Figure 5, Figures 6, 7 and 8). Interestingly, the reduction of ISGs expression in the severe and critical COVID-19 groups of patients coincided with the upregulation of CD55 expression which was also confirmed at the protein level by FACS analysis in each PBMC subpopulation (Figure 5), thus strengthening the validity of our single-cell analysis. The association of CD55 upregulation with reduced ISGs expression identified in monocytes as well as in T and B lymphocytes (Figure) points to a possible involvement of CD55 in the repression of interferon responses signalling displayed in these cellular populations.

CD55 has been linked to the suppression of adaptive immune responses *in-vivo*. In an initial study involving CD55 knock-out mice, CD4^+^ T-cells were reported to excessively produce IFN-γ and IL-2 in response to active immunisation^27^. Subsequently it was shown that lack of CD55 on antigen presenting cells (APCs) and T lymphocytes, during primary T-cell activation, promotes proliferation and increases effector cells^34^. Furthermore, in mouse model studies of CD8^+^ T-cell immune responses to viral infection crossed in CD55 KO background, CD8^+^ T-cell expansion in spleen and lymph nodes was observed alongside an increased number of antigen-specific CD8^+^ T-cells. This expansion resulted in a rapid clearance of viral infection^35^. A subsequent study by the same group demonstrated that treatment of APCs from CD55 deficient mice with an inflammatory stimulus led to aggravated T-cell responses^36^. Finally, CD55 has also been linked to the suppression of NK cell responses ^37^. The association of increased CD55 expression in monocytes T cells and B cells of patients with severe and critical COVID-19 with a reduction of ISGs expression identified in our study is therefore a strong indication of a possible contribution of CD55 in the suppression of type I interferon responses in COVID-19 patients.

Due to the limited number of subjects included in our study, we decided to further examine the validity of our observations by using a large dataset of scRNA-seq analysis previously performed on COVID-19 patients^12^. The study described a comprehensive scRNA-seq analysis of PBMCs from 102 COVID-19 patients. Stephenson et.al utilised the same criteria for patient grouping as those used in our study and included the same groups, except for two additional groups of asymptomatic patients and patients with moderate disease^12^. This facilitated comparisons of our findings for each group and we were thus able to confirm the expression pattern we obtained for CD55 in monocytes and T cells but not for B cells (Figure 2h, Figure 3g and Figure 4f and Supplemental Figure 3). Moreover, the expression observed for the rest of the CRPs were also confirmed suggesting that the discrepancy we observed for CD55 in the B cell population may be due to the smaller number of cells analysed in our study compared to Stephenson et. al., and to variations in the sub clustering methodology and analysis thus resulting in differences in the cell types included and analysed^12^.

We also confirmed the elevated CD55 expression observed in the severe and critical COVID-19 groups of patients at the protein level by FACS analysis. The up-regulated expression was validated in all the PBMC subtypes except for B cells. As anticipated in B cells, upregulation was weaker and only reached statistical significance in the critical COVID-19 group of patients compared to controls (Figure 2). This finding was also in discrepancy to the CD55 expression trend observed using the scRNA-seq dataset of Stephenson et.al.

In summary, in the present scRNA-seq study, we identified a COVID-19 specific expression pattern for complement regulatory protein CD55 characterised by upregulation in monocyte, CD4^+^ and CD8^+^ T cells as well as B cell lineage clusters. Increased expression of CD55 in these subpopulations was associated with decreased type I interferon responses, indicating a potential role of CD55 in COVID-19 severe and acute pathogenesis by promoting suppression of interferon responses. The present study further strengthens the importance of complement activation in COVID-19 and introduces novel molecules of the complement cascade as COVID-19 mediators. Given the diverse array of complement targeted therapeutic strategies our findings may point towards novel therapies in the fight against COVID-19.

## Materials and methods

### Patient samples

Ethical approval was obtained from the Ethics committee of Evangelismos Hospital (approval number: 360, approval date: 17-9-2020, Study title: COVID-19 and Immunological profile). A written informed consent was obtained from all patients or the patient’s next of kin. All patients had a positive polymerase chain reaction (PCR) test for SARS-COV-2 performed on a nasopharyngeal sample. A total of 12 patients were recruited for PBMC isolation for single cell RNA seq analysis. Patients were grouped into those with mild disease (patients admitted to the hospital ward without the need for oxygen supply) (n=4), severe COVID-19 patients (patients who required oxygenation but not ICU admission) (n=4) and critically ill (n=4) (patients who required intubation and ICU admission) (n=4). Healthy individuals (n=4) were used as controls. Blood samples were obtained within 24 hours of admission to either hospital or ICU and processed immediately.

### PBMC isolation for single cell RNA sequencing analysis

PBMC were isolated following a standard protocol. Briefly, 20 ml of whole blood drawn in heparin coated tubes were layered on top of 15 ml of Lymphoprep and separated by gradient centrifugation (700g, 30 min). The central band containing PBMC was extracted (using a sterile Pasteur pipette) into Hanks balanced salt solution (HBSS) (10 ml) and the suspension was centrifuged (700g, 10 min). Cells were counted and processed immediately for scRNA Seq analysis.

### Single cell suspension preparation and generation of droplet-based single cell RNA sequencing data

Four male individual samples per condition were processed (healthy n=4, mild n=4, severe n=4 and critical n=4). Isolated PBMCs were observed and counted with hemocytometer prior to loading. A range of total 106 to 2 × 10^6^ cells were isolated in all 16 samples, and were resuspended to a concentration of 1,000 cells/µl in 0.04% BSA and directly processed for scRNA-seq. Four sample pools were prepared, each aimed to yield 3.000 cells/individual from four individuals (12,000 cells). Each sample pool was loaded into a different lane of a 10× chip (Single Cell G Chip Kit for v3.1). The 10x Chromium controller (10x Genomics), in combination with v3.1 reagents, was used to capture the single cells and generate Single Indexed sequencing libraries, according to the manufacturer’s instructions (document CG000204, RevD). Sequencing was performed with a 150 bp paired-end kit using a custom program (28-8-0-150) on the Illumina NextSeq 500/550 at pMedGR (Medical School, NKUA, Athens, Greece).

### Alignment

Droplet libraries were processed using 10x Genomics Cell Ranger 4.0.0. Reads were aligned to the GRCh38 human genome. The filtered feature matrices were used.

### Doublet Detection

Scrublet ^38^ (v0.2.3 version) was used for the identification of doublets, on each separate sample using the raw counts. Keeping the automatic threshold on the generated doublet score, 8,642 cells were identified as doublets and removed thus ending up with a total of 17,048, 22,997, 7,784 and 16,337 cells for the samples derived from the healthy, mild, severe and critical groups pf patients respectively.

### Quality Control, Preprocessing and Label Transferring

The python package scanpy ^39^ was used for analysis. Combined raw data (64,166 cells) were filtered to remove cells that expressed fewer than 200 genes, genes that were expressed in less than three cells, cells with > 20% mitochondrial reads and cells with > 5,000 n genes by counts. Counts were normalized and log + 1 corrected and highly variable genes (HVGs) were identified. Percentage of mitochondrial reads and the number of total counts were regressed out and the counts were scaled. Using scanpy’s ingest function, that maps labels and embeddings from reference data to query data, we annotated our data on two PBMCs scRNA-seq datasets^13^ that were available at the time, with 647, 366 and 44, 741 cells respectively. Keeping only those cells sharing the same broad annotation in the two reference datasets on the following cell types: monocytes, T cells, B cells, DCs, stem cells, RBCs and platelets for the downstream analysis. The label transferring annotations were confirmed by the expression of known marker genes. As for the cells that the annotation was different between the two different reference datasets, the expression of mixed markers led us to remove them.

### Integration and Sub-clustering

We then subset the three cell types of interest (monocytes, T cells and B cells) and analysed them separately. There was an evident batch effect between the four different conditions, so beginning with raw counts and using the same preprocessing steps we integrated the data using the batch removal tool Batch Balanced KNN ^40^ (BBKNN). Ridge regression for improvement ^41^ of the integration was also performed prior to the final BBKNN step. Clustering was performed using the Leiden algorithm with the resolution being decided based on expression of differentially expressed genes, which were calculated using the Wilcoxon rank sum test. Although doublets were removed in the pre-processing step, clusters from B cells and T cells that expressed mixed markers with other cell types were also removed from further analysis. Annotation of individual clusters was based on literature established marker genes and differentially expressed genes (DEGs). Interferon stimulated genes (ISG) score was calculated using the Scanpy’s tl.score_genes and R package UCell^42^. For the differential expression of CD55 in monocytes, besides the Wilcoxon rank-sum test (scanpy), MAST, DESeq2 and edgeR from R package Libra^43^ were also used.

### Coronascape

Pathway enrichment analysis was performed by Coronascape (https://metascape.org). Upregulated and downregulated DEGs were analyzed, respectively. Pathways with p-value < 0.05 were considered significantly enriched.

### Gene Set Enrichment Analysis (GSEA)

Online at WebGestalt^44^ using the p-value and logFC of the Wilcoxon rank-sum test, we calculated a score by multiplying the -log10 (p-value) with the logFC. Then we ordered the genes in descending order and ran the GSEA using the Hallmark50 Functional Database.

### Flow cytometric analysis for CD55 and CD59 protein expression

For validation of the single cell analysis by FACS, whole blood samples were obtained from patients in the ward with characteristics matching those in the severe COVID-19 group of patients (n=20) (mean age 74.4 ± 12.58) or from the ICU (n=20) (mean age 65.35 ± 21.37). Samples were processed immediately for detection of CD55 and CD59 on monocytes and granulocytes or stored in the -80°C for detection of CD55 and CD59 in CD4^+^ and CD8^+^ T cells and B cells. In order to determine levels of CD55 and CD59 on the various cell types we utilized two separate FACS protocols. A standard protocol used in routine clinical practice for detection of CD55 and CD59 simultaneously in monocytes and granulocytes was used in which 500 µl of freshly drawn blood were added to standard volumes of antibodies (Supplemental Table 1) and incubated in the dark for 15 min. Upon completion of incubation FACS lysing solution (BD Pharmigen) was added and a further incubation for 15 min in the dark was performed. Samples were analysed in the flow cytometer immediately after lysing. The same procedure was utilized for detection of CD55 and CD59 in CD4^+^ and CD8^+^ T cells and B cells with the difference that samples were thawed at room temperature before addition of antibodies. Antibodies used for detection are shown in Supplemental Table 2. The FACS Canto II flow cytometer (BD Biosciences) was used for analysis. Gating strategies are shown in Supplemental Figures 6 and 7. FlowJO software was utilised for analysis of monocyte and granulocyte populations whereas the BD FACSDiva software was used for analysis of the CD4^+^ and CD8^+^ T cells and B cells populations.

### RNA extraction, reverse transcription and Real-time PCR

For analysis of CD55 mRNA levels by Real-time PCR amplification, RNA was extracted from whole blood samples isolated from severe (n=20) and critical (n=20) COVID-19 patients. Whole blood samples were collected in Tempus Blood RNA Tubes (Applied Biosystems) and stored at -80°C. RNA was extracted according to the manufacturer’s instructions. RNA concentration was determined for each sample prior to reverse transcription using NanoDrop One. Reverse transcription reactions were performed using 4 µl of the 5x FastGene® Scriptase II ReadyMix (Nippon Genetics, Duren, Germany) and 100 ng of RNA for each reaction. Reactions were carried out in a CFX90 cycler (BioRad, Hercules, CA) at the following conditions: 25 °C for 10 minutes, 42°C for 60 minutes and 85°C for 5 min. Real time PCR reactions were carried out in a CFX90 cycler. Each reaction consisted of 1 μl primer-probe assay mix (IDT, Coralville, IA), 10 μl Luna Master Mix (Biolabs, Waltham, MA) and 1 μl cDNA. GAPDH was used as a housekeeping gene for data normalization. The sequences for the Real time PCR primers and probes for CD55 mRNA are as follows: forward primer : 5’-CTC ATA TTC CAC AAC AGT ACC GA -3’, reverse primer: TGG TCA GAT ATT GAA GAG TTC TGC -3’, probe: /56-FAM/TGC ATC CCT /ZEN/CAA ACA GCC TTA TAT CAC TC/3IABkFQ/. Reactions were carried out in triplicate and results were analysed by the ΔΔCT method.

### Statistical analysis

Results are reported as absolute numbers, medians, or means and standard deviations, as appropriate. Statistical analysis was performed using the GraphPad Prism 8.0 software for Windows. Data were tested for normality using the Shapiro-Wilks test. Kruskal-Wallis test was used. One-way Anova or Kruskal-Wallis analysis of variance was used in the case of data displaying normality or not respectively the Fisher’s least significant difference (LSD) test was used for post-hoc analysis. Spearman correlation was used for data correlation. Receiver operating characteristic (ROC) analysis was performed using ICU admission as the classification variable and CD55 mRNA levels on admission as prognostic variables. P< 0.05 was considered statistically significant.

## Acknowledgements

We acknowledge financial support of this work by The Greek Research Infrastructure for Personalised Medicine (pMedGR) (MIS 5002802), funded by the Operational Programme “Competitiveness, Entrepreneurship and Innovation” (NSRF 2014-2020) and co-financed by Greece and the European Union (European Regional Development Fund). We also acknowledge support by the Hellenic Foundation for Research and Innovation (H.F.R.I.) under the “1st Call for H.F.R.I. Research Projects to support Faculty members and Researchers and the procurement of high-cost research equipment” (project Single.Out, #3780, to GK).

