## Supplemental Table 2 for "Upregulation of CD55 complement regulator in distinct PBMC subpopulations of COVID-19 patients is associated with suppression of interferon responses"

| Antibody | Fluorochrome | Clone/ Cat no | Manufacturer |
|---|---|---|---|
| Anti-Hu CD45 Pacific Orange | Pacific Orange | 2D1/PO-160-T100 | exbio |
| Anti-Hu CD3 Pacific Blue | Pacific Blue | 10.1/561191 | exbio |
| Anti-Hu CD4 PerCP-Cy | PerCP-Cy | MEM-241 / T9-359-T100 | exbio |
| Anti-Hu CD8 APC-Cy | APC-Cy | MEM-31 / T4-207-T100 | exbio |
| Anti-Hu CD19 PE-Cy | PE-Cy | LT19 / T7-305-T100 | exbio |
| Anti-Hu CD55 | FITC | MEM-118/1F-230-T100 | exbio |
| Anti-Hu CD59 | PE | MEM-43/1F-233-T100 | exbio |

**Supplemental Table 2.** List of antibodies utilized for detection of CD55 and C59 in T and B lymphocytes by flow cytometric analysis.
