## Supplemental Figure 1 for "Upregulation of CD55 complement regulator in distinct PBMC subpopulations of COVID-19 patients is associated with suppression of interferon responses"

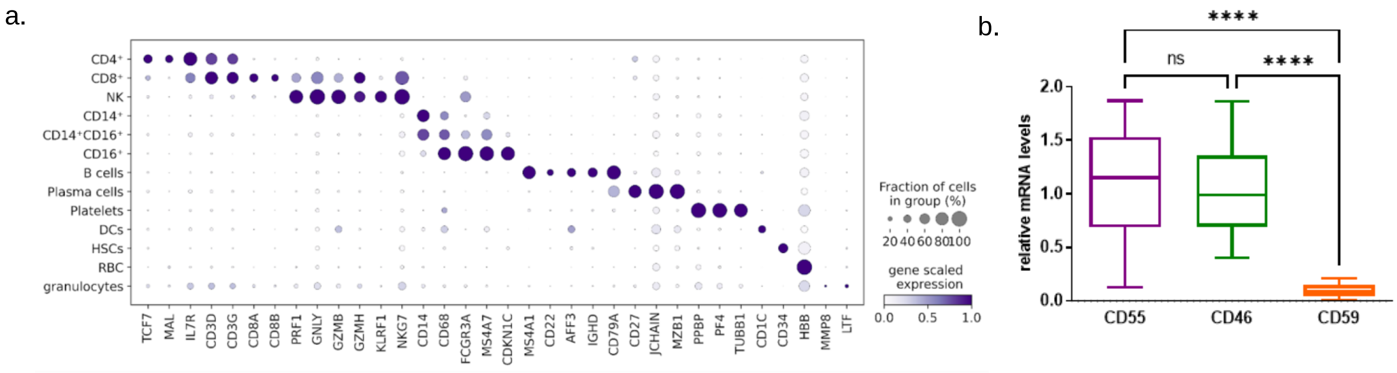


**Supplemental Figure 1. Molecular characterisation of PBMC subpopulations.** a) Dot plot representing expression of established marker genes utilised for PBMC subpopulation identification during integrated data analysis b) Quantitation of CRP expression levels in whole blood samples of healthy individuals by Real-time PCR. One way ANOVA was used for data analysis. Post-hoc analysis was performed by the least significant difference (LSD) test. ****p<0.0001.
