## Supplemental Figure 2 for "Upregulation of CD55 complement regulator in distinct PBMC subpopulations of COVID-19 patients is associated with suppression of interferon responses"

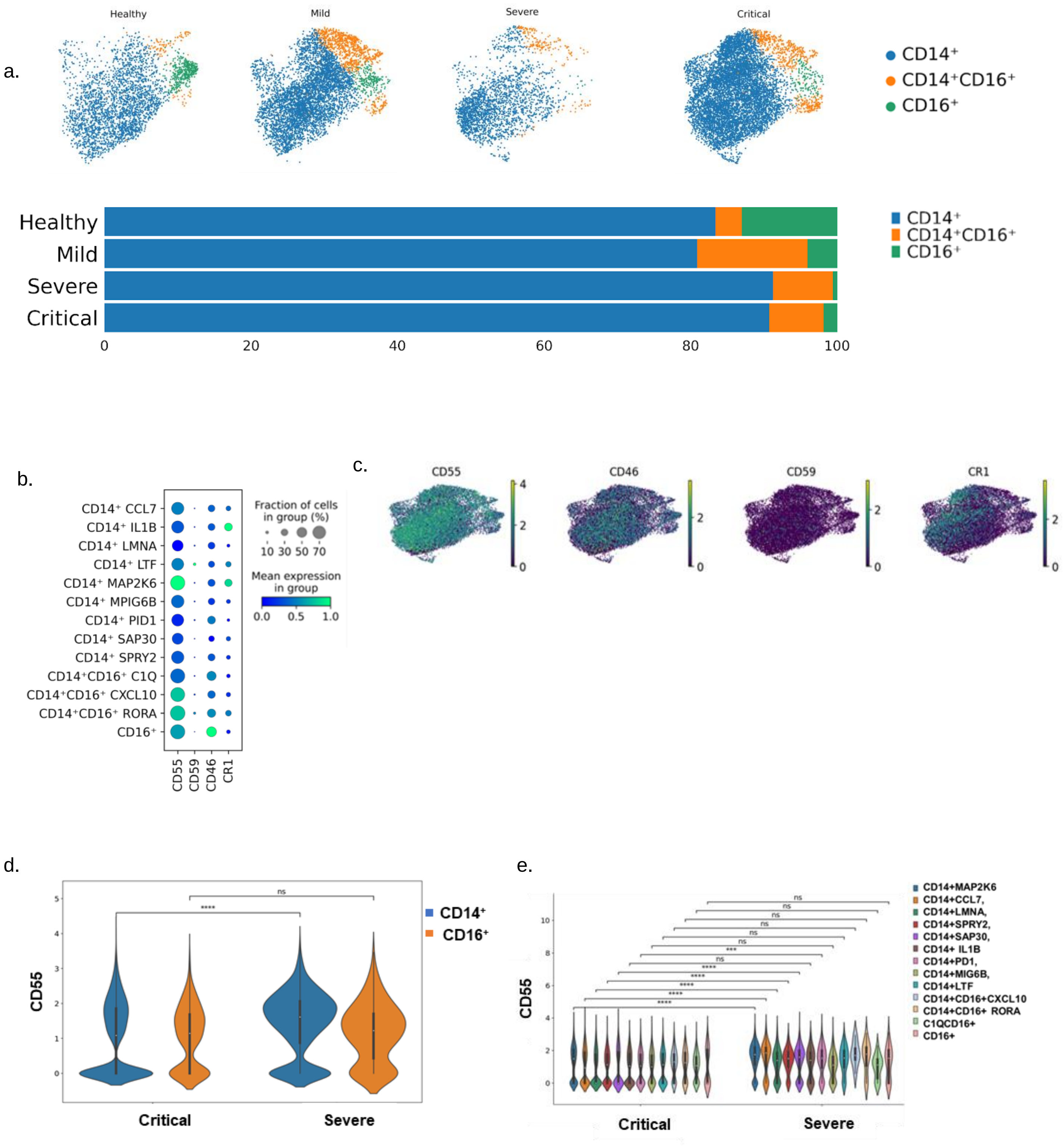


**Supplemental Figure 2. Subclustering analysis of the monocyte subpopulation. a)** CD14⁺ CD16⁺ inferred monocyte-lineages projected in the UMAP space upon integrated analysis in each separate disease state and in healthy controls. b) Dot plot depicting the CRP expression levels in each monocyte subpopulation and c) feature plots of these marker genes d) Violin plots for CD55 gene expression in CD14 and CD16 monocytes in critical versus severe COVID-19 patients e) CD55 expression in monocyte clusters in critical versus severe COVID-19 patients. Statistical analysis performed by the Wilcoxon rank sum test ***p<0.001, ****p<0.0001.
