## Supplemental Figure 3 for "Upregulation of CD55 complement regulator in distinct PBMC subpopulations of COVID-19 patients is associated with suppression of interferon responses"

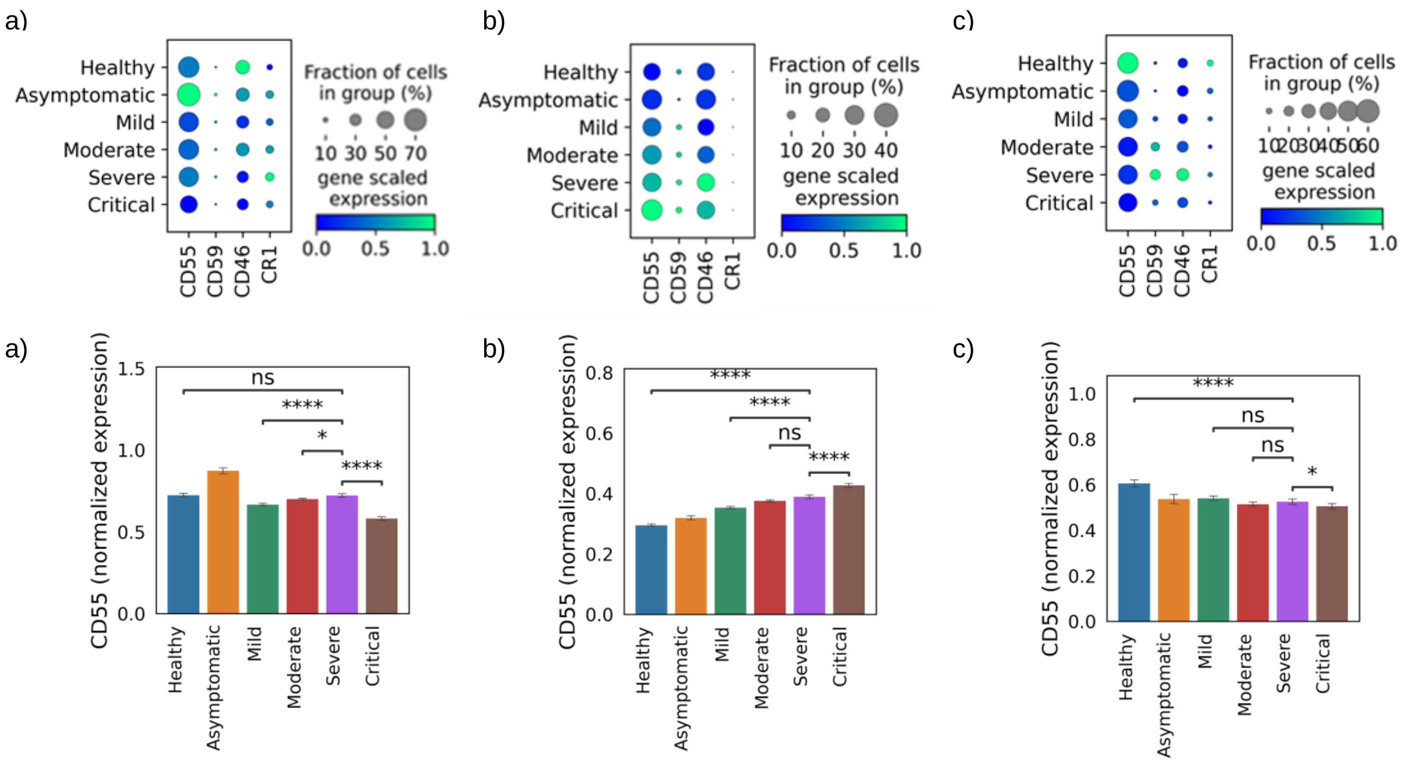


**Supplemental Figure 3. CRP expression patterns by scRNA Seq and flow cytometric analysis**. a) Single cell analysis in a large dataset obtained from deposited data verified the CRP expression patterns observed by our analysis in a) monocytes, b) T cells and c) B cells. Statistical analysis performed by the Wilcoxon rank sum test *p<0.05, ****p<0.0001
