## Supplemental Figure 4 for "Upregulation of CD55 complement regulator in distinct PBMC subpopulations of COVID-19 patients is associated with suppression of interferon responses"

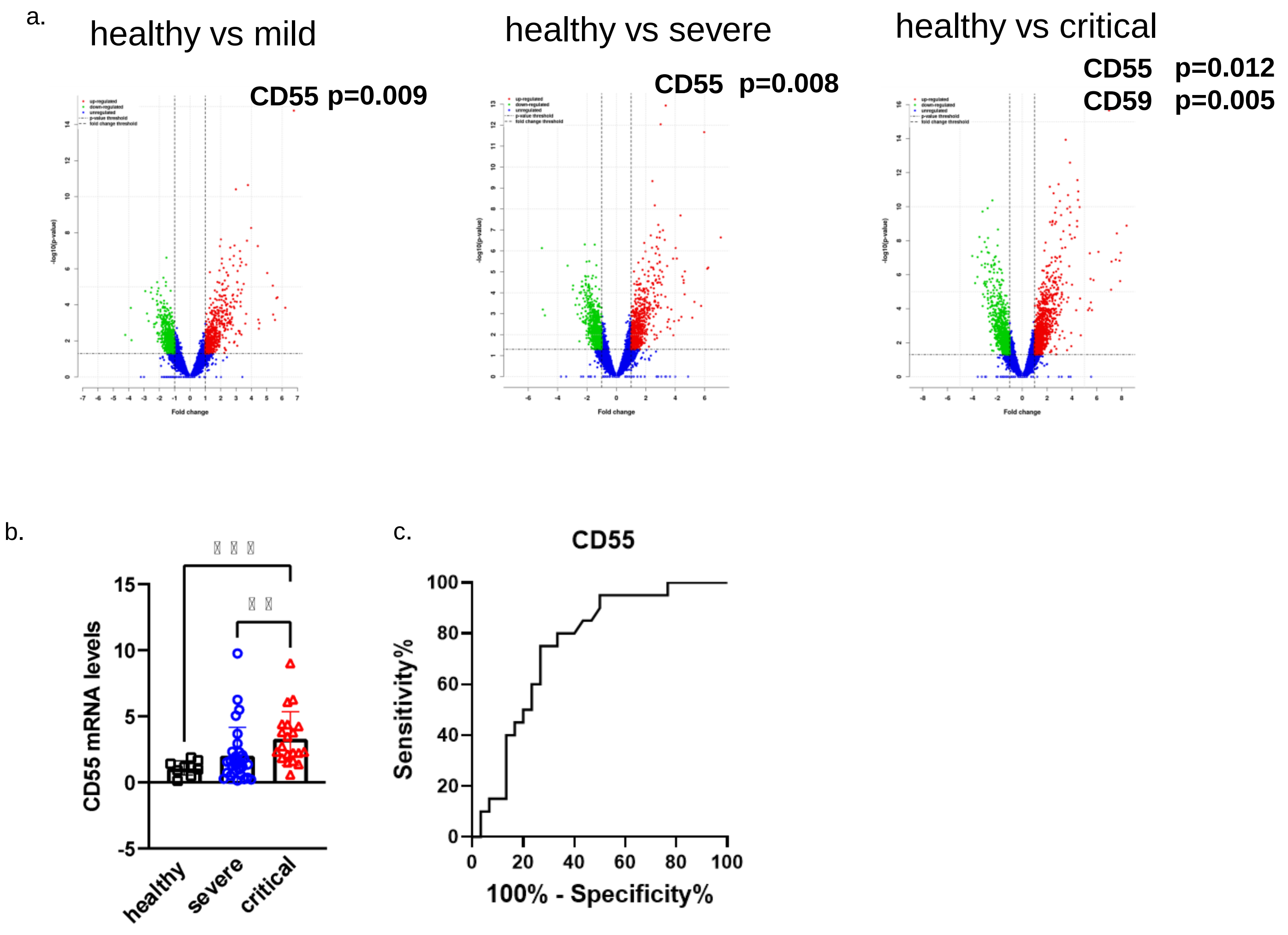


**Supplemental Figure 4. CD55 and CD59 expression levels in COVID-19 patients**. a) CD55 upregulation confirmed by bulk RNA sequencing performed on isolated PBMCs in mild versus healthy (p=0.009), severe COVID-19 patients versus healthy controls (p=0.008) and in critical versus healthy (p=0.012). CD59 upregulation in critical COVID-19 patients versus healthy controls (p=0.005). b) mRNA levels of CD55 in COVID-19 severe and critical patients compared to healthy controls. **p<0.01, ***p<0.001. Statistical analysis was performed by Kruskal-Wallis non parametric test for more than two groups comparisons, the Least significant difference (LSD) test was used for post hoc analysis. c) Receiver operating curve analysis for CD55 for prediction of ICU admission.
