## Supplemental Figure 5 for "Upregulation of CD55 complement regulator in distinct PBMC subpopulations of COVID-19 patients is associated with suppression of interferon responses"

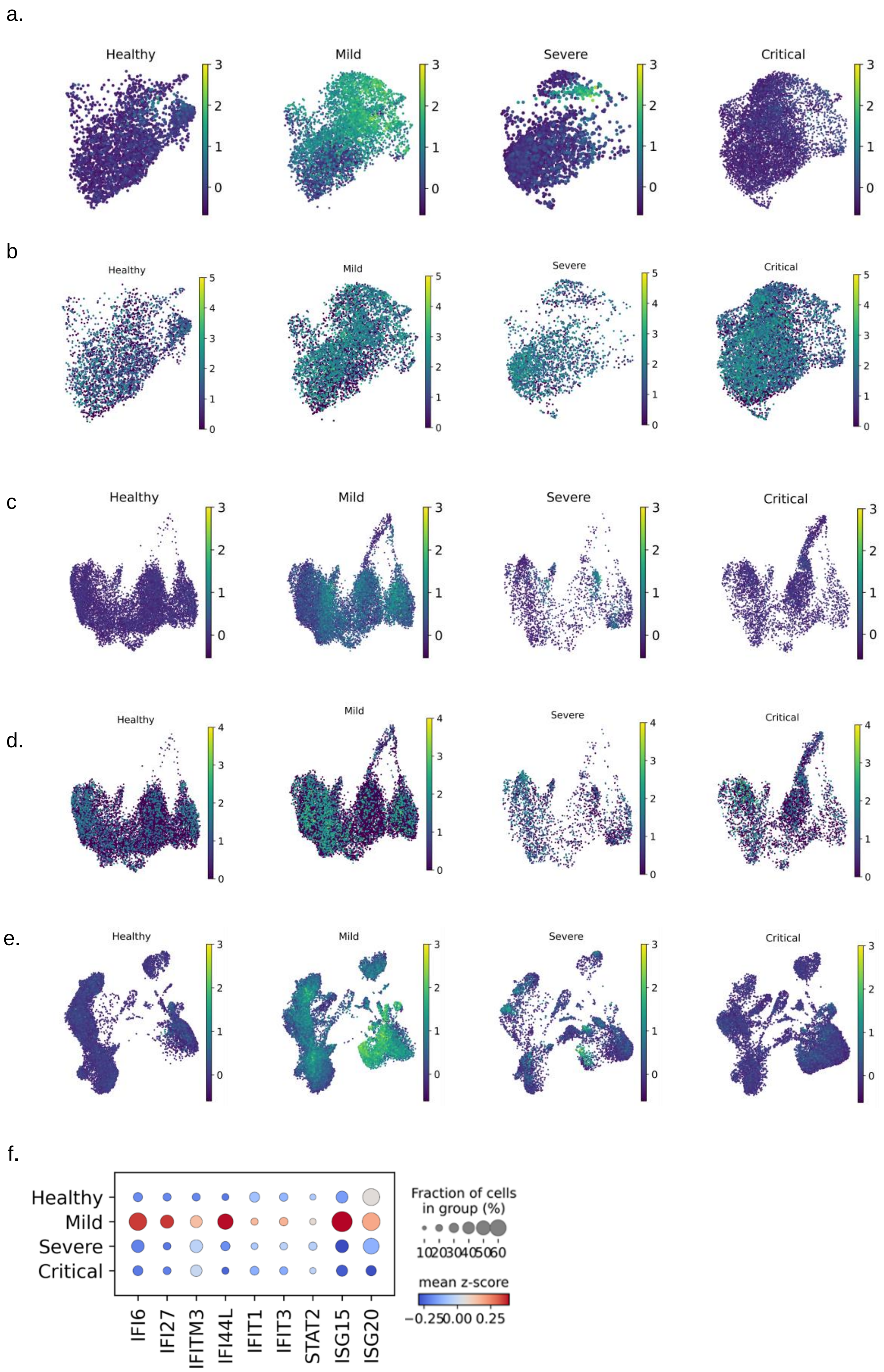


**Supplemental Figure 5. ISG score expression changes in the monocyte, T cell and PBMC population of COVID-19 patients.** Feature plots demonstrating a) ISGs expression on monocyte integrated analysis UMAP, b)CD55 expression on monocyte integrated analysis UMAP, c) ISGs expression on T cell integrated analysis UMAP, d) CD55 expression on T cell integrated analysis UMAP e) ISGs expression on PBMC integrated analysis UMAP and f) dot plot of ISGs score in PBMC of COVID-19 patients and healthy controls.
