## Supplemental Figure 6 for "Upregulation of CD55 complement regulator in distinct PBMC subpopulations of COVID-19 patients is associated with suppression of interferon responses"

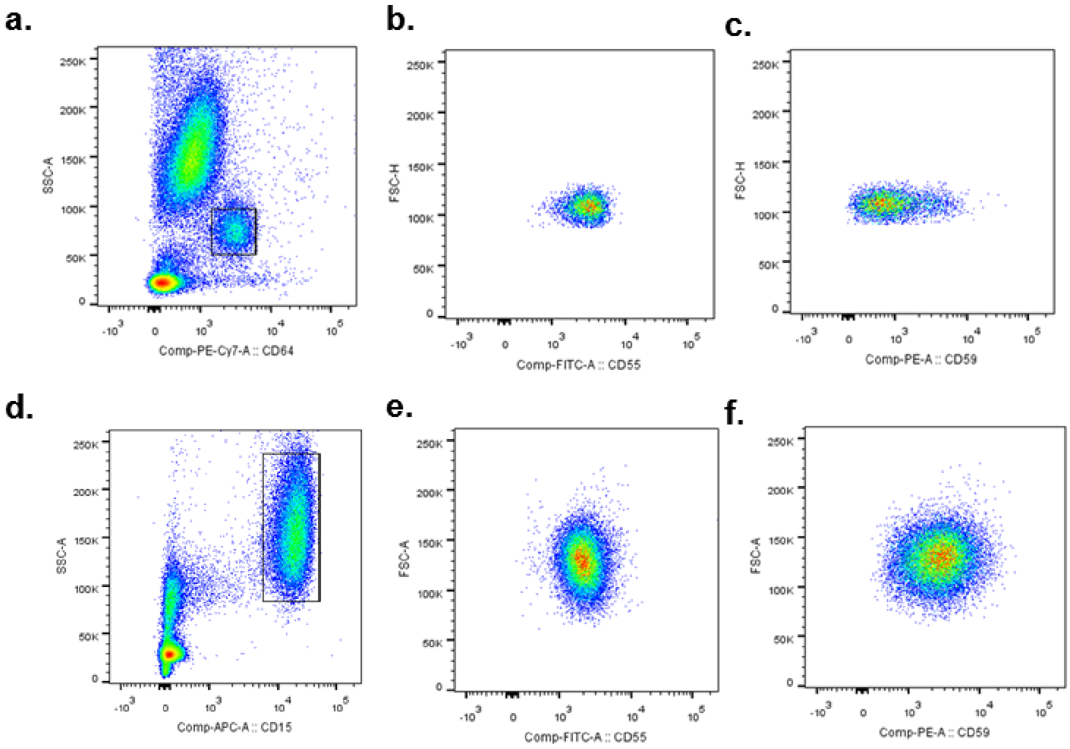


**Supplemental Figure 6. Gating strategy for CD55 and CD59 detection in monocytes and granulocytes by Flow cytometric analysis.** Representative diagram of gating strategy performed using FlowJo software showing a) gated CD64$^+$ cells in order to select for CD45$^+$ for analysis of CD55 (b) and CD59 (c) positive cells within the monocyte population. D) gating of CD15$^+$ cells e) CD55 positive granulocytes and f) CD59 positive granulocytes.
