## Supplemental Figure 7 for "Upregulation of CD55 complement regulator in distinct PBMC subpopulations of COVID-19 patients is associated with suppression of interferon responses"

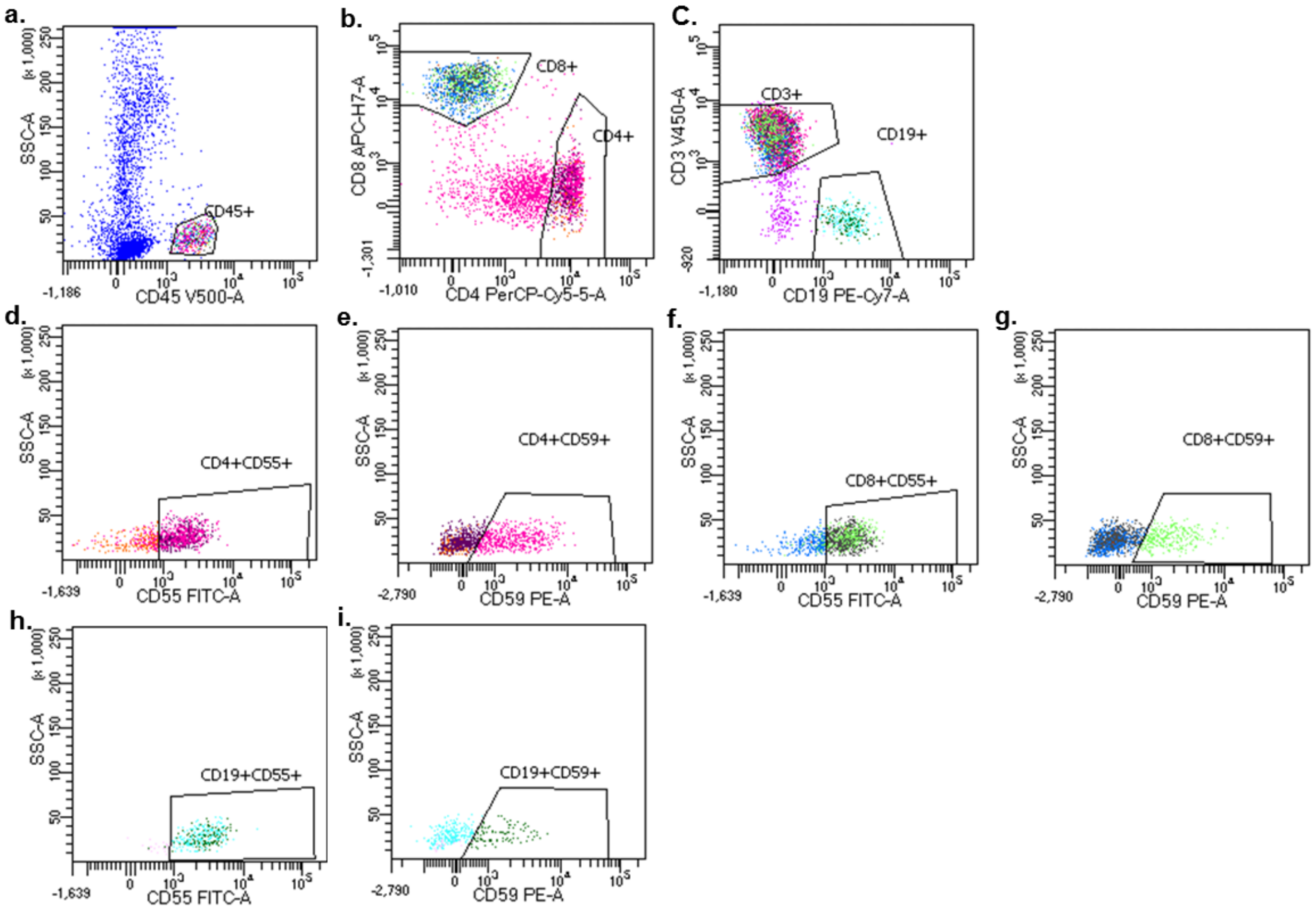


**Supplemental Figure 7. Gating strategy for CD55 and CD59 detection in CD4$^{+}$, CD8$^{+}$ and B cells by Flow cytometric analysis.** Representative diagram of gating strategy using the Diva software of the FACS CANTO flow cytometer showing a) gated CD45$^{+}$ cells in order to select for (b) CD4$^{+}$ and CD8$^{+}$ (c) CD19$^{+}$ cells D) gating of CD4$^{+}$CD55 positive cells e) gating of CD4$^{+}$CD59 positive cells f) ) gating of CD8$^{+}$CD55 positive cells g) gating of CD8$^{+}$CD59 positive cells h) gating of CD19$^{+}$CD55 positive cells i) gating of CD19$^{+}$CD59 positive cells.
