## Supplemental Table 1 for "Upregulation of CD55 complement regulator in distinct PBMC subpopulations of COVID-19 patients is associated with suppression of interferon responses"

| Antibody | Fluorochrome | Clone/ Cat no | Manufacturer |
|---|---|---|---|
| Mouse anti-Human CD45 | APC-H7 | 2D1/560178 | BD Pharmigen |
| Mouse anti-Human CD64 | PC7 | 10.1/561191 | BD Pharmigen |
| anti-human CD15 (SSEA-1) | APC | HI98/301908 | BioLegend |
| Anti-Hu CD55 | FITC | MEM-118/1F-230-T100 | exbio |
| Anti-Hu CD59 | PE | MEM-43/1F-233-T100 | exbio |

**Supplemental Table 1.** List of antibodies utilized for immune profiling of patients by flow cytometric analysis.
